# Quantitative Framework for Assessing Mesenchymal Stem Cell Quality Driven by Poised Enhancer Decommissioning

**DOI:** 10.64898/2026.04.14.718591

**Authors:** Keiko Hiraki-Kamon, Atsushi Wada, Takashi Suyama, Yumi Matsuzaki, Hidemasa Kato

## Abstract

Mesenchymal stem cell (MSC) heterogeneity and conventional phenotypic criteria limitations represent major bottlenecks in therapeutic manufacturing. Here, we present a framework to prospectively identify naturally superior MSCs by shifting from superficial markers to the digital quantification of fundamental epigenetic flaws in inferior clones. We show that intrinsic MSC functional decline is driven by targeted hypermethylation of poised enhancers, causing paradoxical derepression of developmental genes. We term this process poised enhancer decommissioning (PEnD). By isolating this universal decay axis from donor-specific immunological variability, we translate this complex epigenetic state into a streamlined transcriptomic signature: the Poised Enhancer-related Gene Expression (PErGE) score. Overcoming the limitations of standard in vitro differentiation assays, our approach enables accurate, donor-independent prediction of long-term proliferative potential. Together, our findings establish a mechanism-based biomarker of cellular aging, and provide a readily applicable tool to improve the quality control of next-generation MSC-based therapies.

## Introduction

Mesenchymal stem cells (MSCs), also known as mesenchymal stromal cells,^1^ are a cornerstone of regenerative medicine. However, their clinical success is hampered by inconsistent therapeutic outcomes.^2,3^ Conventionally, this inconsistency is attributed to the biological heterogeneity of MSC populations, and their susceptibility to cellular senescence during the extensive in vitro expansion required for clinical use.^4–7^ Yet, emerging single-cell transcriptomic studies suggest that such functional divergence may not be merely an acquired culture artifact but rather rooted in pre-existing cellular states.^8,9^ Current quality control measures, including the binary “pass/fail” International Society for Cell and Gene Therapy (ISCT) criteria and subjective judgment of skilled practitioners, lack the granularity to resolve these inherent differences and predict functional decline.^10–13^ This observation highlights an unmet need for objective, molecular-based metrics that can prospectively identify functionally superior, intrinsically senescence-resistant MSC populations.

Although genetic engineering approaches have shown promise in enhancing MSC therapeutic properties, such as creating senescence-resistant cells with antiaging effects,^14–16^ these approaches face regulatory hurdles and barriers to clinical adoption.^17^ Therefore, identifying and isolating naturally occurring MSC populations that possess inherently superior therapeutic properties without the complexities of genetic modification is urgently required.

In our previous work, we identified such a candidate population enriched for “elite” clones with proliferative capacity, termed rapidly expanding clones (RECs), by isolating MSCs based on NGFR/THY1 coexpression at the single-cell level.^18^ Crucially, by sorting directly from the uncultured bone marrow mononuclear fraction, this strict single-cell isolation functions as an immediate “physical quarantine”, protecting pristine cells from the paracrine senescence signals of their neighbors. However, functional heterogeneity persists even within this strictly isolated population.^19^ Because paracrine confounding factors are eliminated in our system, the persistent functional heterogeneity strongly implies that the divergent fates of these clones, elite versus senescence-prone, are predetermined, thus raising the question of what molecular mechanisms dictate this intrinsic fate.

Epigenetic drift, the time-dependent accumulation of epigenetic alterations within the in vivo niche, is a fundamental driver of cellular aging.^20–22^ This drift involves changes in DNA methylation, an epigenetic mark whose function is strictly context-dependent. Crucially, unlike analog and reversible transcriptional states or labile histone modifications, DNA methylation serves as a highly stable, binary epigenetic record^23,24^, a digital sensor that irreversibly captures transient perturbations in chromatin architecture. Canonically, whereas methylation at promoter regions is a mechanism for gene silencing, methylation within gene bodies is positively correlated with active transcription, ensuring transcriptional fidelity.^23–26^ In MSCs, epigenetic drift disrupts this delicate regulatory landscape. Although such epigenetic decay is conventionally dismissed as a mere artifact of in vitro expansion,^22,27–31^ whether these flaws are actually preexisting states that inherently dictate cell fate remains unclear. By shifting our focus from defining ideal MSC traits to quantifying these permanent epigenetic signatures, we aimed to uncover the fundamental drivers of inherent cell quality.

Therefore, we undertook epigenetic and transcriptomic analyses to identify the preexisting molecular mechanisms defining the functional potential of MSCs. Specifically, we investigated whether the epigenetic integrity of poised enhancers, developmental regulatory elements that should remain silent yet reversible to preserve future lineage options, is the decisive factor in dictating MSC quality. We aimed to translate this mechanistic understanding into a predictive molecular signature. Critically, we sought a framework that could disentangle intrinsic cellular potency from donor-specific variability, thereby enabling robust, prospective selection of elite MSCs based on their preestablished endogenous properties.

## Results

### Isolation and characterization of distinct MSC clonal populations from human bone marrow

By analyzing single-cell–derived, isogenic MSC clones with divergent long-term proliferative capacities, we were able to examine intrinsic determinants of MSC functional quality independently of donor variability and genetic manipulation. We performed single-cell sorting of MSCs from uncultured human bone marrow aspirates based on NGFR^+^/THY1^+^ coexpression. This dual-positive population represents an extremely rare fraction (typically <0.1% of bone marrow mononuclear cells). Furthermore, among these sorted cells, only a minor subset (approximately 10 clones per 96-well plate) achieved the rapid initial proliferation required for subculture within 2 weeks. We selectively isolated these highly proliferative elite clones, which we termed RECs (Figure 1A).^18^ However, we observed that extended culture unmasked a profound functional heterogeneity within this strictly isolated REC population. Based on their long-term proliferative potential, we classified these clones into two subtypes: “true” RECs (tRECs), which maintained their robust proliferative capacity, and subrapidly expanding clones (SrECs), which exhibited premature growth deceleration. Despite being functionally relevant, this classification highlighted the reliance on subjective, experience-dependent judgments for assessing MSC quality as a bottleneck in the field. Therefore, we aimed to establish objective, molecular-based metrics to overcome this limitation.

**Figure 1.**
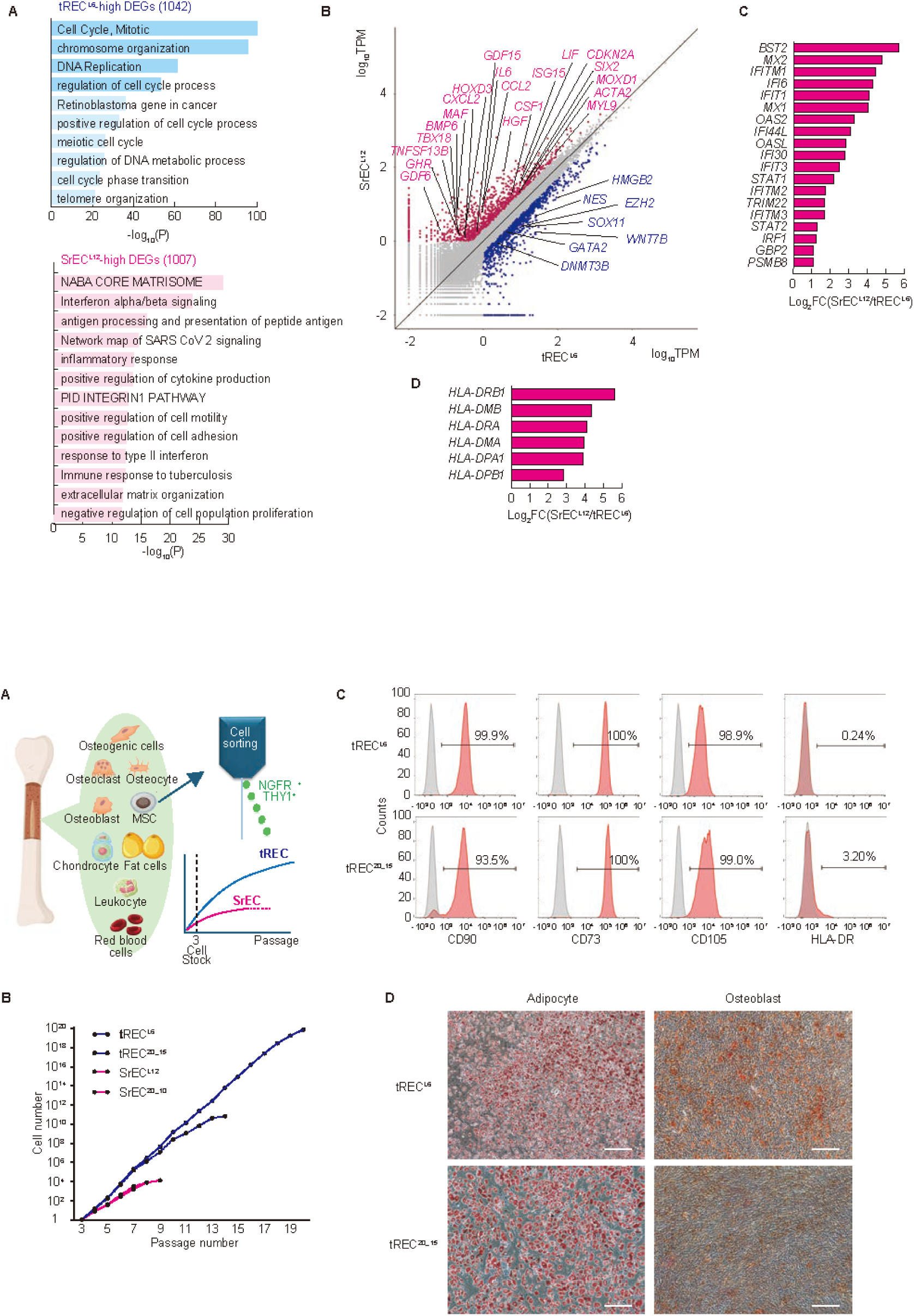
Isolation and functional characterization of tREC and SrEC. (A) Schematic of the isolation procedure for MSC clones based on NGFR^+^/THY1^+^ coexpression, and their subsequent functional classification into tRECs and SrECs based on long-term proliferative potential. (B) Long-term proliferation assay of four single-cell-derived MSC clones. tREC clones (tREC^L6^, tREC^20_15^) exhibit sustained, long-term expansion, whereas SrEC clones (SrEC^L12^, SrEC^20_10^) undergo premature growth arrest. The elite clone tREC^L6^ achieved a total expansion of >10^26^-fold from a single cell, equivalent to 86 cell doublings. (C) Flow cytometry analysis confirming that representative tRECs at passage 5 (P5) meet the minimal ISCT criteria: positive for CD73, CD90, and CD105, whereas negative for HLA-DR. (D) Representative images showing that tRECs retain multilineage differentiation potential into osteogenic (Alizarin Red S) and adipogenic (Oil Red O) lineages. Scale bar = 200 μm. Abbreviations: MSC, mesenchymal stem cells; tRECs, “true” rapidly expanding clones; SrECs, subrapidly expanding clones

To investigate the molecular basis of this heterogeneity, we selected two pairs of isogenic clones (tREC^L6^/SrEC^L12^ and tREC^20_15^/SrEC^20_10^) from independent donors for analysis (Supplementary Table S1). Among these, tREC^L6^ emerged as an exemplar of MSC function. Serial passaging analysis revealed its exceptional proliferative capacity, achieving a cell increase of at least 10^26^-fold (equivalent to 86 cell doublings) from a single cell, one of the highest expansion potentials recorded for nonengineered human MSCs (Figure 1B). Despite this massive expansion, the tREC^L6^ clone met the defined ISCT criteria⁹ (Figure 1C). Although the other highly proliferative clone (tREC^20_15^) exhibited minor deviations in surface marker expression (CD90 = 93.5%; HLA-DR = 3.2%), both tREC clones retained adipogenic and osteogenic differentiation potentials (Figure 1D). This naturally occurring senescence resistance established tREC^L6^ as a benchmark for defining the molecular hallmarks of baseline MSC potency.

### Transcriptome analysis reveals bifurcation into proliferative and senescent states

We conducted comparative transcriptome analysis to define the molecular signatures underlying these functional states and identified a stark contrast between the two populations (Supplementary Figure S1A). Gene ontology (GO) analysis showed that consistent with their proliferative phenotype, tRECs were enriched for cell cycle and DNA replication pathways. Notably, tRECs expressed higher levels of *EZH2*, the catalytic subunit of the Polycomb Repressive Complex 2 (PRC2) recently identified as a driver of cellular rejuvenation,^32^ and the antiaging factor *HMGB2*.^33^ Conversely, SrECs were characterized by an aging-associated signature, with elevated expression of the cell cycle inhibitor *CDKN2A*^34^ and enrichment of extracellular matrix (“matrisome”)-^35^ and lineage differentiation-associated genes (Supplementary Figure S1B).

To dissect these divergent signatures, we focused our analysis on the isogenic pair tREC^L6^ and SrEC^L12^, which exhibited a pronounced phenotypic difference. This comparison revealed that SrEC^L12^ had acquired an inflammatory and aging phenotype. In particular, SrEC^L12^ demonstrated a senescence-associated secretory phenotype (SASP), marked by the upregulation of factors such as *IL6* and *GDF15*,^36^ alongside enrichment for genes involved in interferon signaling and MHC class II antigen presentation (Figure 2). By contrast, tREC^L6^ maintained an antiaging transcriptional profile, characterized by high expression of factors such as *EZH2*, *GATA2*^37^ and *HMGB2*. Furthermore, the tREC^L6^ transcriptome was enriched for neural crest-associated markers (*SOX11*, *NES*),^38,39^ suggesting that its properties may be linked to a primitive developmental origin. This early hint of lineage-specific signatures suggested that the functional divergence between tRECs and SrECs might reflect distinct developmental roots rather than stochastic culture-associated degradation. These coordinated transcriptional shifts prompted us to investigate whether an underlying epigenetic mechanism stabilizes these divergent cell states.

**Figure 2.**
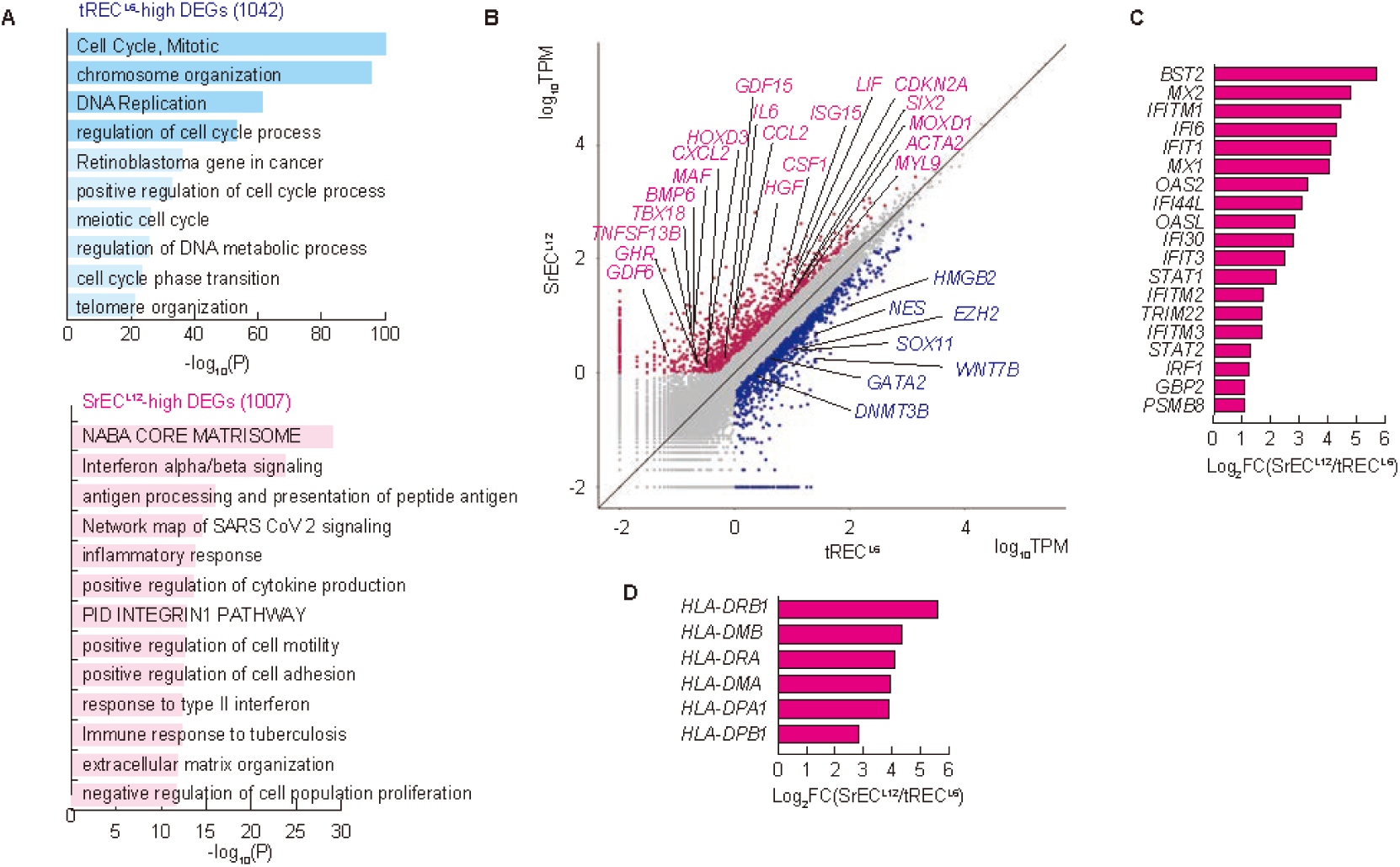
Transcriptomic signatures distinguish the potent tREC state from senescent SrEC state. (A) GO analysis of genes highly expressed in tREC^L6^ (tREC-high DEGs) and SrEC^L12^ (SrEC-high DEGs). tREC-high DEGs are enriched for proliferation-related pathways (e.g., cell cycle, DNA replication), whereas SrEC-high DEGs show enrichment for pathways related to the matrisome, interferon signaling, and antigen presentation. (B) Scatter plot of RNA-seq data comparing the transcriptomes of tREC^L6^ (*x*-axis, *log*_10_ TPM) and SrEC^L12^ (*y*-axis, *log*_10_ TPM). Genes associated with the potent tREC state (e.g., *EZH2*, *HMGB2*; blue dots) and senescent SrEC state (e.g., *CDKN2A*, *IL6*; red dots) are highlighted. (C and D) Bar plots quantifying the significant upregulation (*Log*_2_ fold change) of interferon-responsive genes (C) and MHC class II HLA genes (D) in SrEC^L12^ compared with those in tREC^L6^, indicating an acquired proinflammatory and immunogenic phenotype. Data represent the fold change between these two representative isogenic clones (n = 1) derived from RNA-seq analysis. Abbreviations: DEG, differentially expressed genes; GO, Gene Ontology; tRECs, “true” rapidly expanding clones; SrECs, subrapidly expanding clones; TPM, transcripts per million

### SrEC functional decline is driven by targeted DNA hypermethylation of poised enhancers

To identify the epigenetic basis of this transcriptional divergence, we performed whole-genome bisulfite sequencing. Typically, culture-induced cellular senescence is characterized by a stochastic, global loss of DNA methylation (epigenetic drift).^28,30,31^ However, the critical epigenetic changes defining the SrEC state were not driven by this canonical global hypomethylation (Supplementary Figure S1C). Instead, our analysis revealed that the divergence was dictated by highly targeted changes at specific genomic loci.

Indeed, analysis of differentially methylated regions (DMRs) revealed a clear dichotomy: tREC^L6^-hypermethylated DMRs were broadly scattered across intergenic regions whereas SrEC^L12^-hypermethylated DMRs were selectively concentrated in cis-regulatory elements, especially enhancers (Figure 3A, Supplementary Figure S2A-D). To understand the reason for the targeting of these enhancers, we examined their chromatin state. We found that the SrEC^L12^-hypermethylated enhancers were enriched for the repressive histone marker H3K27me3 (Figure 3B, C). This marker defines a class of regulatory elements known as “poised enhancers”,^40^ which are kept silent but primed for activation during development.

**Figure 3.**
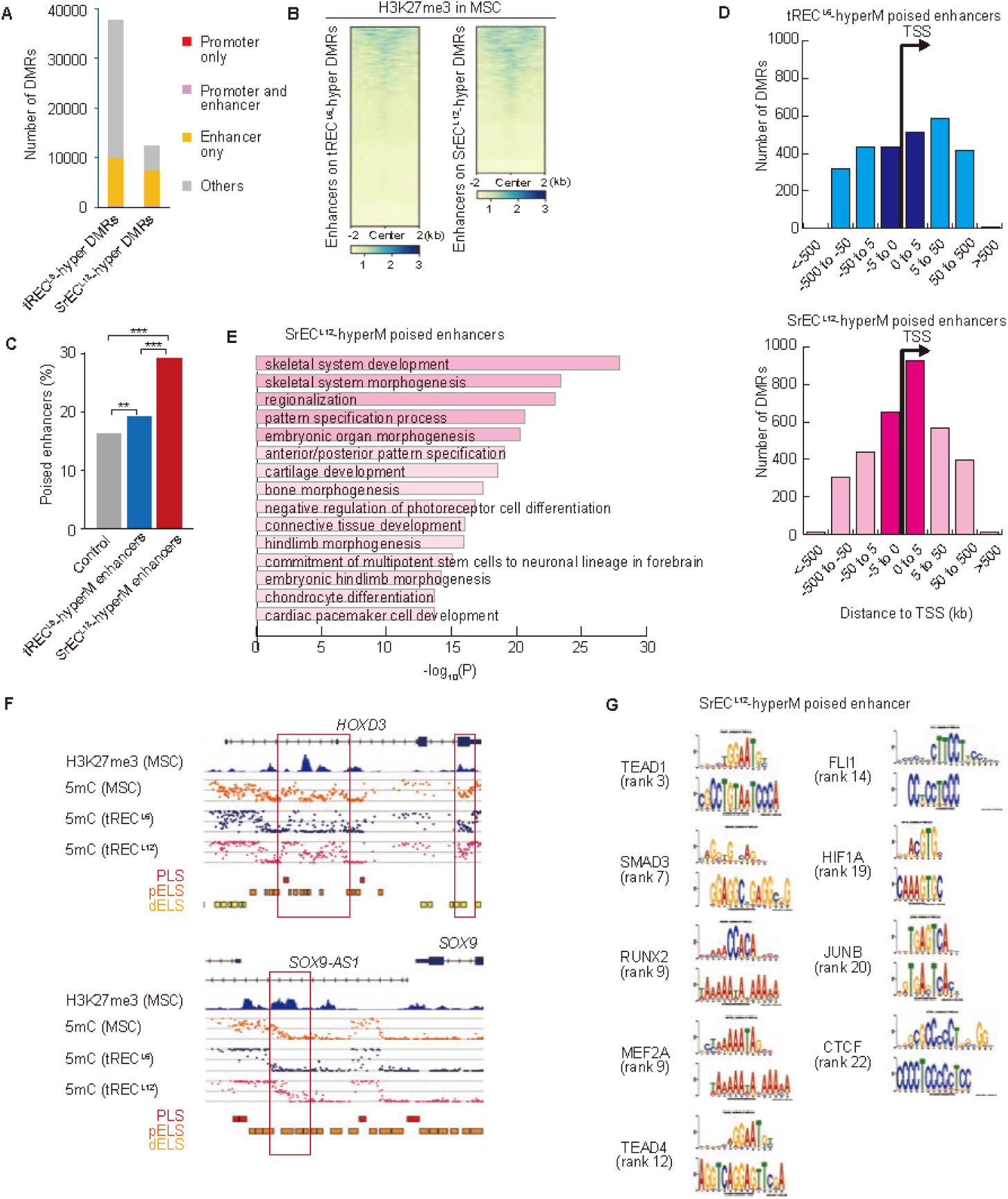
SrEC functional decline is driven by the targeted DNA hypermethylation of poised enhancers. (A) Genomic distribution of DMRs. SrEC^L12^-hypermethylated DMRs are selectively enriched in enhancer regions compared with those in tREC^L6^-hyperM DMRs. (B and C) Heatmap (B) and quantification (C) demonstrating that SrEC^L12^-hyperM enhancers are strongly enriched for the repressive histone marker H3K27me3 in MSCs. This colocalization defines them as “poised enhancers”. (D) Analysis of genomic location relative to the transcription start site (TSS). Although tREC^L6^-hypermethylated poised enhancers display a broad and relatively even distribution, those in SrEC^L12^ are characteristically concentrated immediately downstream of the TSS. (E) GO analysis of genes associated (within 1000 kb) with SrEC^L12^-hyperM poised enhancers. (F) Representative genome browser tracks for key developmental regulators (*HOXD3*, *SOX9*). These loci illustrate the “poised enhancer decommissioning (PEnD)” mechanism wherein the gain of DNA methylation (5mC) in SrEC^L12^ (red track) occurs at the H3K27me3-marked (blue track) poised enhancers. (G) HOMER motif analysis shows that SrEC^L12^-hyperM poised enhancers are enriched for the binding sites of key mesenchymal transcription factors (e.g., RUNX2, SMAD3), which are expressed in SrECs (see Supplementary Table S2). Abbreviations: DMR, differentially methylated regions; GO, Gene Ontology; MSC, mesenchymal stem cells; tRECs, “true” rapidly expanding clones; SrECs, subrapidly expanding clones; TPM, transcripts per million

Notably, these SrEC^L12^-hypermethylated poised enhancers were concentrated immediately downstream of the transcription start sites (0–5 kb), locating them directly within the proximal gene body (Figure 3D). Given this specific spatial distribution, we next asked which functional gene categories are associated with these epigenetic alterations.

### Hypermethylated poised enhancers target skeletal development pathways

GO analysis revealed that the SrEC^L12^-hypermethylated poised enhancers are associated with genes involved in skeletal system development, that is, the canonical differentiation fate of MSCs (Figure 3E, Supplementary Figure S2E). These include developmental regulators such as the Polycomb target *HOXD3*^41^ and the master chondrogenic factor *SOX9*^42^ (Figure 3F). As established previously, DNA methylation at these enhancer sites displaces the repressive PRC2 complex, thus removing the epigenetic barrier that normally keeps them silent. To determine whether these now-accessible enhancers could indeed be activated, we investigated the second crucial component: the presence of the necessary transcription factors (TFs).

Motif analysis revealed that these enhancers were enriched for the binding sites of master mesenchymal regulators, including RUNX2^43^ and SMAD3^44^ (Figure 3G). RNA-seq data further confirmed that these TFs were expressed in SrECs (Supplementary Table S2). This finding completed the overall picture, wherein the epigenetic “lock” (PRC2) on developmental enhancers in SrECs is broken by DNA methylation, whereas the transcriptional “keys” (TFs) required for activation are already present, providing a molecular basis for their divergent fate and lineage restriction.

The contrasting methylation patterns highlight a fundamental aspect of functional MSCs: the preservation of poised chromatin states that maintain developmental plasticity. SrEC^L12^ exhibited slower cell proliferation and demonstrated aging signatures, coinciding with a restrictive epigenetic landscape driven by the hypermethylation of enhancers essential for future MSC lineage differentiation (Figure 3F). Together, these findings indicate that the targeted hypermethylation of poised enhancers represents not a passive, culture-induced epigenetic drift but a specific loss of regulatory potential, a process we refer to as poised enhancer decommissioning (PEnD).

### PEnD drives transcriptional changes through PRC2 displacement and promotes SrEC differentiation

We then examined the transcriptional consequences of aberrant DNA methylation at poised enhancers in SrEC^L12^ cells, to translate the loss of poised regulatory states into coordinated changes in gene expression and MSC fate. Differential expression analysis of the 1654 genes associated with SrEC^L12^-hypermethylated poised enhancers identified 100 upregulated and 117 downregulated genes (Figure 4A). This bidirectional effect revealed a dual regulatory consequence of PEnD (Figure 4B).

**Figure 4.**
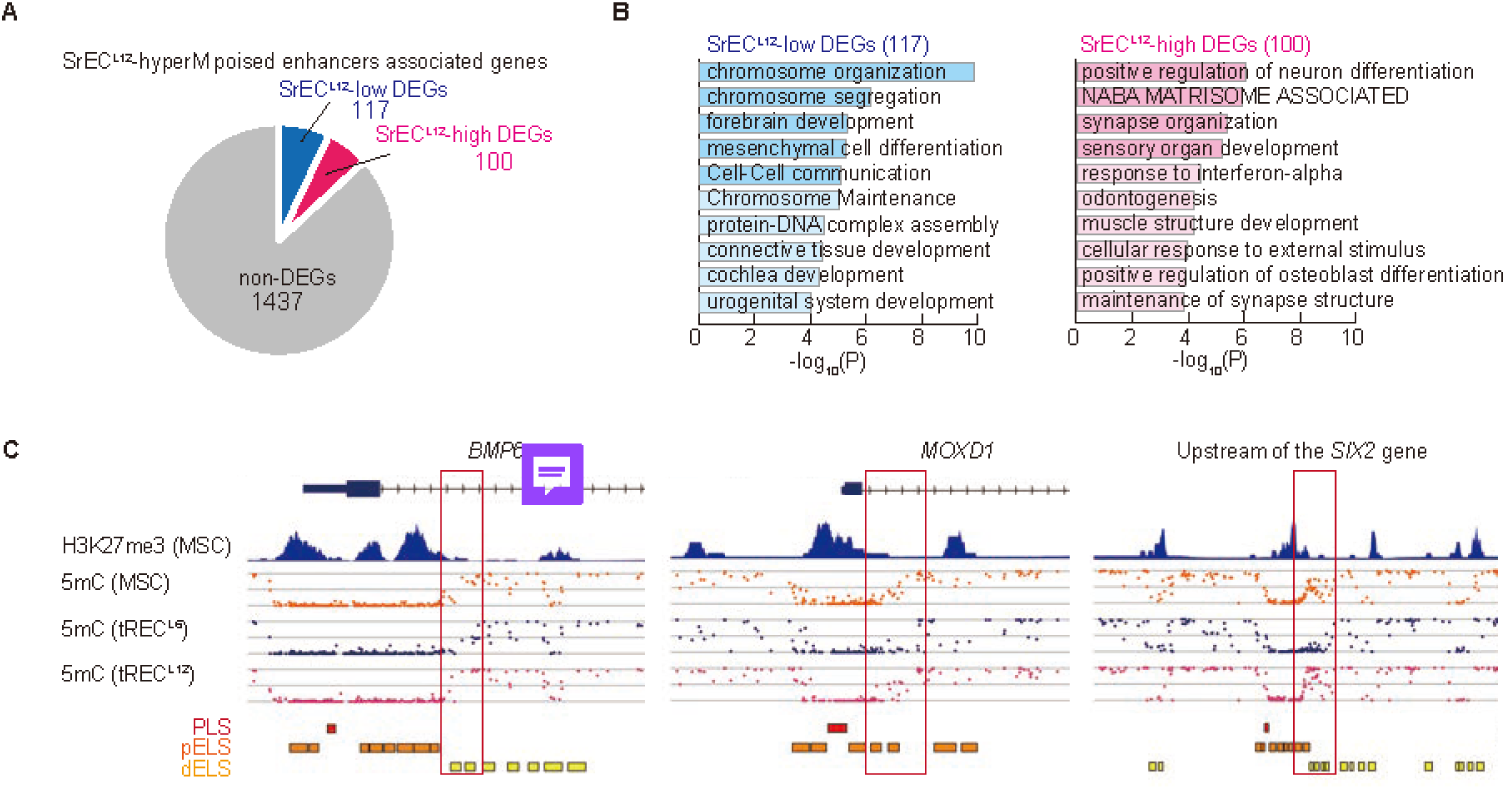
Hypermethylation of poised enhancers drives a dual transcriptional program of functional decline. (A) Pie chart categorizing the 1654 genes associated with SrEC^L12^-hyperM poised enhancers. Differential expression analysis identified 100 upregulated (SrEC^L12^-high DEGs) and 117 downregulated (SrEC^L12^-low DEGs) genes, demonstrating a dual regulatory effect. (B) GO analysis of the DEG sets identified in (A). Downregulated genes (SrEC^L12^-low) are associated with maintaining mesenchymal identity (e.g., mesenchymal cell differentiation), whereas upregulated (derepressed) genes (SrEC^L12^-high) are linked to lineage-specific differentiation programs (e.g., positive regulation of neuron differentiation, NABA MATRISOME ASSOCIATED). (C) Representative genome browser tracks for key developmental genes (*BMP6*, *MOXD1*, *SIX2*) paradoxically derepressed in SrEC^L^^12^. Abbreviations: DMR, differentially methylated regions; GO, Gene Ontology; tRECs, “true” rapidly expanding clones; SrECs, subrapidly expanding clones

The downregulation of a subset of these genes fits the canonical role of DNA methylation, which typically represses transcription by inhibiting transcription factor binding, and recruiting histone deacetylases.^45^ However, at a specific subset of the targeted poised enhancers, this hypermethylation triggers PEnD, leading to a paradoxical outcome: the activation of normally silenced genes. This derepression is consistent with a mechanism wherein DNA methylation antagonizes PRC2 binding and function on silent enhancers.^46,47^ The eviction of the PRC2 complex and subsequent loss of the H3K27me3 mark lead to the derepression of developmental genes previously maintained in a poised, silent state.

Indeed, in SrEC^L12^ cells, this PEnD-induced PRC2 displacement mechanism accounts for the unscheduled upregulation of lineage-specific differentiation genes that should remain silenced in multipotent MSCs. The upregulated gene set was enriched for differentiation-related factors, including *DLX1* (4.8-fold increase), *DLX2* (3.5-fold increase),^48^ and *CDON* (3.0-fold increase).^49^ Furthermore, the SrEC^L12^-upregulated differentially expressed genes (DEGs) included interferon response genes and skeletal lineage determinants, such as *BMP6*, *MOXD1*, and *SIX2* (Figure 4C).

These expression patterns indicate that, unlike pristine tREC^L6^ cells, SrEC^L12^ cells are locked in a predifferentiated state primarily attributed to the PEnD-mediated derepression of lineage-specific genes. This mechanistic insight highlights that aberrant DNA methylation restricts the multipotent state not merely through canonical gene silencing but crucially by triggering PEnD, which unleashes unscheduled differentiation programs.

### A poised enhancer gene signature prospectively predicts MSC functional potential

To quantitatively project the functional divergence driven by PEnD into a reduced transcriptional space, we performed principal component analysis (PCA) following log-transformation with postcorrection zero restoration (see Methods for details). When PCA was applied to the global transcriptome (all genes), the first principal component (PC1) predominantly captured donor-to-donor variability rather than the functional cellular state (Figure 5A, right). However, when we restricted the unsupervised analysis strictly to the 1654 PEnD-targeted poised enhancer-associated genes, the mathematical maximization of variance completely shifted. In this PEnD-focused PCA, PC1 (accounting for 23.09% of the variance) effectively overcame donor-specific backgrounds and completely segregated pristine tRECs from deteriorated SrECs based solely on their functional state (Figure 5A, left).

**Figure 5.**
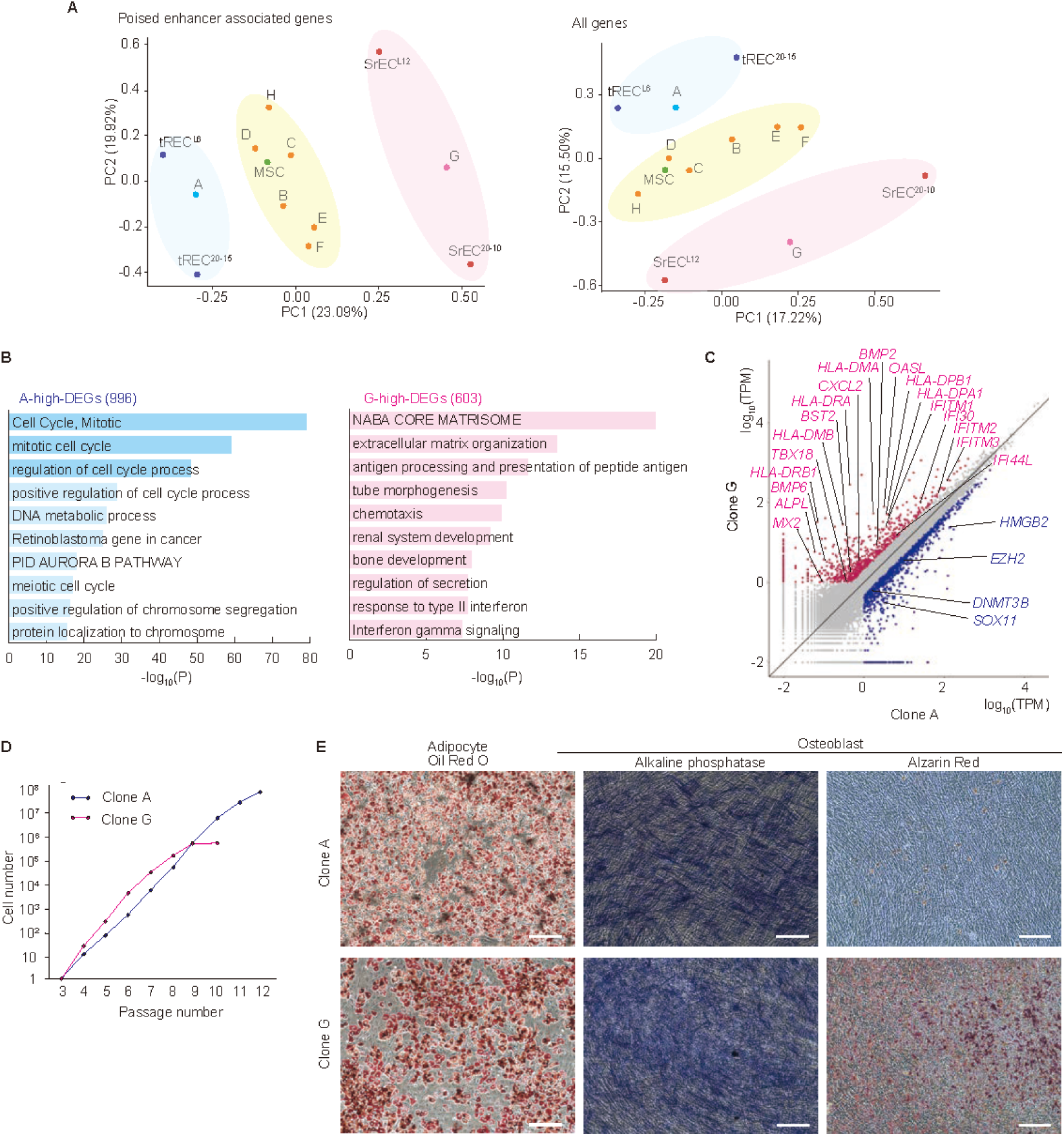
A poised enhancer signature prospectively predicts MSC functional potential. (A) PCA of MSC clones. The poised enhancer-associated gene set (left) provides superior functional classification compared with the whole transcriptome (right). PC1, representing the primary axis of functional quality, effectively segregates high-quality tRECs (negative PC1) from low-quality SrECs (positive PC1). The model prospectively classified clone A as a tREC and clone G as a SrEC. (B and C) Transcriptomic analysis confirming the PCA classification. GO analysis (B) shows that clone A demonstrates a proliferative signature (e.g., cell cycle, mitotic), whereas clone G exhibits a prodifferentiation signature (e.g., NABA CORE MATRISOME, bone development). The differential gene expression scatter plot (C) highlights key genes, including the early osteoblast marker *ALPL*, which is highly expressed in clone G. (D) Long-term proliferation assay functionally validating the prediction. Clone A (predicted tREC) demonstrates vastly superior expansion capacity, surpassing clone G (predicted SrEC) by more than two orders of magnitude. (E) Multilineage differentiation of clones A and G. Despite both clones retaining adipogenic (Oil Red O) and osteogenic (alkaline phosphatase, Alizarin Red S) potential, clone G exhibited enhanced late-stage osteogenesis (Alizarin Red S staining), consistent with its molecular signature indicating a predifferentiated state. Scale bar = 200 μm. Abbreviations: GO, Gene Ontology; MSC, mesenchymal stem cells; PCA, principal component analysis; tRECs, “true” rapidly expanding clones; SrECs, subrapidly expanding clones

As expected from this segregation, PC1 high-loading genes, which indicate strong contributions to this principal component, were highly enriched for multiple adverse traits, including differentiation, fibrosis, senescence, and immunogenicity (Figure 2). tRECs consistently occupied the negative side of this axis, indicating that the separation is driven by a coordinated signature of genes expressed at lower baseline levels in tRECs but inappropriately upregulated in SrECs. This pattern reflects the PEnD mechanism: the methylation-mediated derepression of developmental genes that should remain strictly silenced in potent stem cells.

Crucially, the isolation of PC1 demonstrated that SrEC characteristics are not random artifacts of in vitro expansion but rather represent an intrinsic axis of predetermined cellular fate arising from PEnD. This axis spans from a repressed, potent state (negative loadings) to a derepressed, differentiated state (positive loadings) and is exemplified by the early osteoblast marker *ALPL*^50^ and the myofibroblast lineage regulator *NOTCH3*^51^ being among the top positive-loading genes. Notably, *NOTCH3* upregulation coincided with the expression of contractile cytoskeleton genes, such as *ACTA2* and *MYL9*, whereas the stemness guardian *NOTCH1*^52^ was conversely associated with the tREC state (Supplementary Table S3). Consequently, unsupervised analysis independently extracted a biological signature reflecting both differentiation and physical remodeling.

By contrast, PC2 primarily segregated the clones by donor. Gene set enrichment analysis (GSEA) revealed that this axis captures variable immune landscapes (e.g., interferon response), likely reflecting donor-specific physiological states (Supplementary Figure S3B). Crucially, our analysis successfully isolated the robust, donor-independent quality metric of PC1 from these extrinsic sources of variability.

### Prospective validation confirms the predictive power of the poised enhancer signature

To prospectively validate our framework, we analyzed eight new candidate MSC clones. The PCA-based model unequivocally classified clone A as a high-quality tREC and clone G as a low-quality SrEC (Figure 5A). Their respective transcriptomes substantiated this classification; clone A was enriched for cell cycle genes, whereas clone G displayed the characteristic SrEC signature of high extracellular matrix- and immune response-related gene expression (Figure 5B, C). Indeed, the prodifferentiation signature of clone G significantly overlapped with that of SrEC^L12^, confirming a conserved molecular program of intrinsic lineage restriction across different donors (Supplementary Figure S4).

We then subjected these clones to a head-to-head functional challenge. The results of a long-term serial passaging assay decisively confirmed our prediction: clone A ultimately surpassed clone G by over two orders of magnitude, validating the predictive power of the framework (Figure 5D). Furthermore, although both clones retained differentiation potential, clone G exhibited enhanced late-stage osteogenesis (Figure 5E), consistent with its molecular signature indicating a predifferentiated state. Importantly, the superior proliferative capacity of tREC-like cells was not associated with malignant transformation, as they lacked *TERT* expression^53^ while maintaining robust expression of key tumor suppressors (*TP53*, *RB1*) and a network of genome integrity-related genes (Supplementary Figure S5). Together, these results indicate that MSC functional quality can be understood as a continuum defined by the inherent integrity of poised regulatory states, rather than as a binary property inferred from surface markers or short-term functional assays.

## Discussion

The therapeutic paradigm for MSCs is shifting from a focus on differentiation potential to properties such as senescence resistance.^16^ Recent landmark single-cell studies have highlighted the profound population heterogeneity that complicates this goal, identifying the proliferative apex within the native niche^8^ and suggesting that functional divergence may be rooted in preexisting cellular states. Building on these transcriptomic snapshots, our deep epigenetic profiling of isogenic REC clones provides a mechanistic basis for this inherent heterogeneity. Our findings strongly suggest that the irreversible functional restriction of MSCs is dictated by deep-seated epigenetic differences present from the outset. Uncovering this, however, required a fundamental reorientation in the means by which MSC quality is evaluated, a shift from cataloging cellular states to identifying the initial, irreversible epigenetic events that underpin them.

Conventional approaches to MSC quality assessment rely on defining positive markers of potency, a strategy inherently blind to preexisting epigenetic flaws. We inverted this logic. By isolating naturally occurring isogenic clones, we did not presuppose a specific axis of variation; rather, we allowed the interclonal heterogeneity to reveal the structure of the problem. The identification of tREC^L6^, a clone with near-zero transcriptional noise at developmental loci and exceptional expansion potential, provided the essential empirical anchor. Against this pristine reference, the transcriptomic “noise” in inferior clones was no longer a nuisance to be filtered out but the underlying biological signal to be decoded. This epistemological shift revealed that poised enhancer decommissioning (PEnD) is a fundamental epigenetic mechanism underlying intrinsic MSC functional restriction.

To fully appreciate this mechanism, the concept of the poised enhancer needs to be reconsidered in the context of adult stem cell biology. Although poised enhancers were first described as markers of developmental potential in embryonic stem cells, our findings indicate that they serve a distinct, and critical role in adult stem cells: preserving a latent regulatory architecture that safeguards long-term multipotency.^40^ Our work identified the targeted hypermethylation of poised enhancers, and the resulting “decommissioning” of this regulatory blueprint, as a critical epigenetic feature that defines naturally occurring, lineage-restricted subpopulations within the bone marrow. Despite appearing to be essential for physiological tissue homeostasis, this inherent heterogeneity presents a challenge for regenerative medicine, which relies on the isolation of highly potent, naive stem cells.

In SrECs, the derepression of developmental genes coincides with targeted DNA methylation at poised enhancers. This pattern is consistent with the well-established mutually antagonistic relationship between DNA methylation and Polycomb repression,^46,47^ in which the presence of 5mC is known to antagonize the repressive PRC2 complex. Unlike analog RNA expression changes, the presence of 5mC at these loci constitutes a stable, binary record. Crucially, in a fully potent tREC, DNA methylation at these PRC2-bound enhancers is maintained at a “zero baseline.” Against this absolute zero, the natural presence of 5mC in specific subpopulations represents a categorical divergence, detectable by WGBS with an exceptionally high signal-to-noise ratio. This digital readout effectively captures the preexisting lineage bias associated with attenuated Polycomb repression. Importantly, a recent independent study demonstrating that EZH2 enhancement rejuvenates MSCs provides strong orthogonal support for our PEnD model, reinforcing the premise that PRC2 dysregulation is intrinsically linked to functional deterioration.^32^ Together, these findings strongly support the conclusion that the epigenetic integrity of poised enhancers is a key determinant of the naive multipotent state, distinguishing elite therapeutic candidates from their naturally lineage-biased counterparts.

Building on this epigenetic profile, our findings resolve a critical paradox regarding the context-dependent role of DNA methylation. As established in the Introduction, whereas promoter methylation canonically silences genes, gene body methylation is positively correlated with active transcription. Crucially, our analysis revealed that the poised enhancers targeted by PEnD are spatially concentrated directly within proximal gene bodies (Figure 3D). This specific spatial arrangement suggests a plausible model to explain the remarkable stability of this intrinsic lineage bias during extensive in vitro expansion. Once a developmental gene is derepressed in specific subpopulations, active transcription facilitates the deposition of gene body methylation, a process mediated by transcription-coupled epigenetic modifiers such as the *SETD2*-*DNMT3B* axis.^23–26^ We propose that this transcription-coupled gene body methylation serves as the definitive molecular “lock”. Because this intragenic DNA methylation antagonizes PRC2 binding, the gene would not become resilenced, effectively ensuring transcriptional fidelity and perpetuating the inherent lineage trajectory of the cell. Although direct dynamic validation of this loop in our MSC clones is a subject for future studies, this model effectively bridges the spatial distribution of PEnD with the canonical function of gene body methylation, thereby explaining the maintenance of the preexisting, naturally occurring SrEC state across numerous cellular divisions.

Importantly, the convergence of this self-reinforcing lock-in mechanism with the demonstrated rejuvenating potential of EZH2 reinforcement^32^ carries direct implications for the future experimental approach of the causal validation of PEnD. Because PEnD is not an isolated event at a discrete genomic locus but rather a manifestation of attenuated global PRC2 homeostasis, locus-specific epigenome editing strategies, such as targeted methylation or demethylation of individual enhancers, would be insufficient to establish or refute causality. Instead, meaningful functional interrogation will require approaches that modulate PRC2 activity in a graded, quantitative manner, for example, through degron-mediated titration of EZH2 protein levels. Such experiments, however, must be guided by a prior understanding of the prevalence and consistency of PEnD across diverse donor backgrounds. By establishing the conceptual and analytical framework for detecting PEnD, the present study provides the necessary foundation upon which such a mechanistic dissection can be rationally designed.

The ability to identify this pristine epigenetic state, and to observe its intrinsic lock-in mechanism, was critically dependent on our two-stage NGFR^+^/THY1^+^ single-cell isolation strategy. As independently supported by recent single-cell atlases,^8^ gating on NGFR captures the broad epigenetic spectrum of the niche, whereas THY1 enriches for its proliferative apex. The subsequent clonal expansion from a single cell constitutes, in effect, a physical quarantine. Crucially, this does not negate the reality of culture-induced deterioration; rather, our model delineates its origins. In conventional bulk cultures, cells harboring preexisting lineage biases prematurely undergo PEnD and rapidly exhibit a senescence-associated secretory phenotype (SASP), secreting proinflammatory and profibrotic factors such as IL6, CXCL2, and BMP6. In such a setting, bulk expansion would systematically expose pristine elite cells to this paracrine pollution, potentially triggering a bystander effect that accelerates the secondary, culture-induced fibrotic decommissioning of still-healthy neighbors. By establishing a strict quarantine immediately upon isolation, conceptually akin to a prophylactic senolytic intervention, our single-cell approach successfully decoupled these two phases of decay. This eliminated the confounding noise of paracrine amplification, allowing us to uncover that the fundamental role of intrinsic PEnD events in MSC functional restriction.

Importantly, the primary axis of variation (PC1) derived from our unsupervised analysis of PEnD-associated genes was not engineered by hand-picking markers but emerged naturally from the data. We noted that this specific principal component captured the coalescence of differentiation, fibrosis, and senescence signatures. The emergence of this unified axis provides strong evidence that PEnD is not a bystander but a primary driver of the intrinsic functional divergence observed in MSC populations.

Our transcriptomic analysis revealed a distinctive alteration in Notch signaling components, most notably the prominent expression of *NOTCH3* in SrECs. Although Notch signaling regulates stem cell fate in a context-dependent manner, NOTCH3 is a known driver of myofibroblast differentiation.^54^ Significantly, this *NOTCH3* induction was accompanied by the upregulation of contractile cytoskeleton genes, including *ACTA2* (α-SMA) and *MYL9*.^55^ This gene expression profile resembles the “fibrotic-MSC” or “myofibroblast signature” observed in recent single-cell atlases of the bone marrow niche. These findings suggest that what we conventionally term “senescence” in expanded MSCs may, in part, reflect a preformed “lineage drift” already present in the bone marrow, an intrinsic, clone-specific bias toward a stiff, myofibroblast-like transcriptional state. Rather than representing a stochastic functional decline, our data suggest that MSC heterogeneity comprises a spectrum of physiologically distinct subpopulations, in which certain lineages, although essential for niche homeostasis, possess inherently limited expansion potential for therapeutic applications.

Crucially, our PEnD-focused analysis overcame a fundamental hurdle in MSC quality assessment: the confounding effect of donor-specific immunological signatures. Although global transcriptomic analyses often capture variable immune landscapes reflecting the physiological state or systemic inflammation of the donor,^56,57^ we found that such extrinsic factors are primarily sequestered into a secondary axis of variation (PC2). Indeed, we observed distinct clustering on the PC2 axis that largely mirrored the donor-specific baselines of the original bone marrow lots. Conversely, the primary axis defined strictly by PEnD-associated genes effectively disentangles this donor-specific noise from the intrinsic epigenetic restriction (PC1). By isolating the universal PEnD-related signal from these context-dependent immunological signatures, our framework provides a robust, donor-independent metric that captures the fundamental baseline of stemness across diverse cell lots.

Historically, MSC quality has been evaluated based on the ISCT criteria, which rely on defined surface markers, and tri-lineage in vitro differentiation potential.^11^ However, our data demonstrate that whereas the pristine tREC^L6^ clone perfectly meets these criteria, other highly proliferative, and functionally multipotent clones (such as tREC^20_15^) can exhibit minor deviations from canonical surface markers. Furthermore, recent developmental insights have fundamentally challenged the biological rationale of generic multilineage assays. Nguyen et al. utilized iPSC-derived models to demonstrate that cells with a somitic positional identity intrinsically undergo hypertrophic differentiation and dystrophic calcification, in stark contrast to cranial neural crest cells that form stable hyaline cartilage.^58^ Given that adult bone marrow MSCs are primarily of somitic origin, their physiological fate does not include stable hyaline cartilage formation. To date, the mechanism by which MSCs undergo apparent chondrogenesis in vitro remains undetermined. Rothstein et al. recently provided a compelling mechanistic explanation:^59^ prolonged exposure to TGF-β, a ubiquitous component of standard chondrogenic induction media, can forcibly reprogram restricted cells to acquire cranial-like skeletogenic plasticity. These findings suggest that conventional in vitro chondrogenesis assays may not primarily reflect the intrinsic baseline potency of MSCs but rather their susceptibility to artificial lineage reprogramming driven by nonphysiological cytokines. Together, these discrepancies highlight the profound limitations of relying on superficial phenotypic markers and in vitro differentiation assays that lack a developmental rationale.

Ultimately, to overcome the limitations of these conventional criteria, a fundamental shift in quality assessment is required. Although the fundamental driver of this intrinsic restriction is the epigenetic decommissioning of poised enhancers (PEnD), routine WGBS analysis in clinical manufacturing is highly impractical. By successfully isolating this universal decay axis (PC1) from confounding immunological noise (PC2), we translated this complex epigenetic state into a streamlined transcriptomic signature: the Poised Enhancer-related Gene Expression (PErGE) score. Unlike conventional markers that rely on transient positive indicators, PErGE directly captures the downstream consequences of PEnD, that is, the paradoxical derepression of developmental genes. By monitoring this fundamental biological signal, PErGE offers a predictive, mechanism-based quality control tool that robustly forecasts long-term performance across independent donors. This framework not only advances our mechanistic understanding of intrinsic MSC aging but also provides a practical solution for prospectively identifying naturally occurring elite MSCs toward improving the quality control of next-generation regenerative therapies.

Although our study established a robust framework using multiple isogenic MSC clones, extending this approach to larger cohorts with diverse donor backgrounds will be essential to further define the general applicability of the PErGE signature. Furthermore, despite the PErGE signature successfully predicting the expansion and stemness potential of MSCs in vitro, the present study did not evaluate the therapeutic efficacy of these prospectively identified cells in vivo. Whether the epigenetically “soft” and “poised” state of elite tRECs directly translates to superior tissue regeneration remains a critical question for future studies. Finally, we established a robust correlation between PEnD and functional divergence but did not include direct gain- or loss-of-function experiments to formally demonstrate causality. As PEnD reflects a systemic attenuation of PRC2 homeostasis rather than a defect at any single locus, appropriate causal interrogation will require graded modulation of global Polycomb activity, an experimental strategy that should be informed by large-scale validation studies.

## Resource availability

### Lead contact

Requests for further information and resources should be directed to and will be fulfilled by the lead contact, Hidemasa Kato.

### Materials availability

The specific single-cell-derived MSC clones characterized in the present study are available from the lead contact upon reasonable request. However, owing to the finite nature of these nonimmortalized primary cell stocks, distribution is strictly subject to availability, and will require a completed Material Transfer Agreement (MTA).

### Data and code availability

All raw data for RNA-seq, CUT&Tag, and WGBS have been deposited in the DNA Data Bank of Japan (DDBJ) under the accession numbers DRR665263-DRR665282 and are publicly available as of the date of publication.

This paper does not report any original code.

Any additional information required to reanalyze the data reported in this paper is available from the lead contact upon request.

## Methods

### Selection and culture of rapidly-expanding clones (RECs)

RECs were isolated by integrating high-resolution flow cytometry with a functional screening based on initial growth kinetics.^60^ Fresh whole bone marrow aspirates (Cat#2M-125C, Lonza Japan, Ltd., Tokyo, Japan; Cat#ABM009F, ALCELLS, Alameda, CA, USA;) were processed through a Ficoll-gradient to obtain bone marrow mononuclear cells, which were then incubated with mouse anti-human CD90-APC (Cat#559869, BD Biosciences, Franklin Lakes, NJ, USA) and mouse anti-human CD271-PE (Cat#12-9400, Thermo Fisher Scientific, Waltham, MA, USA). CD90/CD271-double-positive cells (typically representing <0.1% of total mononuclear cells) were collected into 96-well plates using a JSAN cell sorter (BayBioscience, Kobe, Japan).

Cells were cultured in maintenance medium consisting of low-glucose DMEM (FUJIFILM Wako Pure Chemical Corporation, Osaka, Japan), 20% FBS (Cytiva HyClone, Marlborough, MA, USA), 20 ng/mL bFGF (KAKEN PHARMACEUTICAL Co., Ltd., Tokyo, Japan), 0.01 M HEPES (Thermo Fisher Scientific), and 1% penicillin/streptomycin (Meiji Seika Pharma, Tokyo, Japan). REC selection was strictly based on their initial proliferative vigor: only the clones reaching subconfluency within the first 14 d (typically ∼10 clones per 96-well plate, or ∼10% of sorted cells) were harvested and passaged into 6-well plates (P1). Clones failing to achieve rapid expansion during the initial 2-week period were excluded. Subsequently, cells were expanded into T75 flasks (P2), and cryopreserved as P3 stocks in STEM-CELLBANKER (ZENOGEN PHARMA, Fukushima, Japan) at -80 °C. These P3 stocks were utilized for differentiation, and serial-passaging assays. Cells were maintained at 37 °C and 5% CO_2_, with medium changes conducted every 3 d. P5 subconfluent cells were used for RNA and DNA extraction.

### Serial passaging analysis

To evaluate long-term proliferative potential, P3-stocked cells were thawed, and serially passaged as previously described.^60^ Cells were seeded at a density of 2.0 × 10^5^ cells per 100-mm dish (approximately 2500 cells/cm^2^), expanded to 80-90% confluence, then harvested, and replated. Cell numbers were tracked at each passage to calculate cumulative expansion factors. Total proliferative capacity was calculated as the product of initial single-cell-to-P3 expansion (approximately 10^7^-fold) and subsequent P3-to-endpoint expansion.

### Initial empirical classification of MSC clones

Initial classification of MSC clones into tREC and SrEC categories was performed by skilled cell culture practitioners with extensive MSC culture expertise based on empirical observations of growth kinetics, morphological changes, and passage-dependent proliferative decline. Despite being functionally relevant, this subjective assessment exemplified the nonstandardized nature of current MSC quality evaluation practices, and motivated us to develop objective, molecular-based classification criteria.

### MSC differentiation

#### Adipocyte differentiation

Adipogenic differentiation was performed based on a published protocol, with modifications.^61^ Trypsin-harvested cells were cultured for 2 weeks in adipogenic differentiation medium consisting of DMEM, 20% FBS, 0.01 M HEPES, 1% penicillin/streptomycin, 0.2 mM indomethacin (Wako), 0.5 mM isobutylmethylxanthine (Nacalai Tesque, Kyoto, Japan), and 1 μM dexamethasone (Wako). Differentiated cells were stained with Oil Red O (Muto Pure Chemicals, Tokyo, Japan).

#### Osteogenic differentiation

For osteogenic differentiation, a previously published protocol was employed, with modifications.^62,63^ Harvested MSCs were cultured for 3 weeks in osteogenic differentiation medium containing DMEM, 20% FBS, 0.01 M HEPES, 1% penicillin/streptomycin, 10 mM β-glycerophosphate (Merck, Darmstadt, Germany), 50 μM L-ascorbic acid phosphate (Wako), and 10 nM dexamethasone (Wako). Differentiated cells were stained with Alizarin Red S (Muto Pure Chemicals) for assessing calcium deposition, and alkaline phosphatase activity was evaluated using an ALP Staining Kit (Takara Bio Inc., Kusatsu, Japan) according to the manufacturer’s instructions.

### Flow cytometry analysis

Cells were suspended in PBS containing 2% fetal bovine serum (FBS) at a concentration of 2 × 10^6^ cells/mL and stained for 30 min on ice using the following antibodies: mouse anti-human CD73-PE (Cat#344003, BioLegend, San Diego, CA, USA), CD90-APC (Cat#559869, BD Biosciences), CD105-APC (Cat#323208, BioLegend), and HLA-DR-FITC (Cat#IM1638U, Beckman Coulter). Propidium iodide (BD Biosciences) was used to exclude dead cells. Analyses were performed on a CytoFLEX (Beckman Coulter, Brea, CA, USA), and data were processed using FlowJo software (BD Biosciences).

### RNA sequencing and analysis

#### Library preparation and sequencing

Total RNA was purified using the RNeasy Plus Mini Kit (QIAGEN, Hilden, Germany), following the manufacturer’s instructions. Library preparation, and sequencing were performed by Genome-Lead Co., Ltd. (Kagawa, Japan). Ribosomal RNA was removed using the NEBNext rRNA Depletion Kit v2 (Human/Mouse/Rat) (New England Biolabs, Ipswich, MA, USA), and RNA-seq libraries were prepared using the Illumina Stranded mRNA Prep, Ligation kit (Illumina, San Diego, CA, USA). Libraries were sequenced on an Illumina NovaSeq 6000 platform, generating 151-base paired-end reads.

#### Data analysis

RNA-seq reads were aligned to the human reference genome (GRCh38/hg38) using STAR software with default parameters. Transcripts per million mapped reads (TPM) were calculated using RSEM with default parameters. Differentially expressed genes (DEGs) between REC and SrEC populations were identified using the R package DESeq2 with cutoffs of absolute log_2_(fold change) >1 and Benjamini-Hochberg adjusted P-value <0.05 to control false discovery rate (FDR). Because pairwise comparisons between individual isogenic clones (tREC^L6^ vs. SrEC^L12^; clone A vs. clone G) lack biological replicates and thus preclude statistical modeling by DESeq2, DEGs for these comparisons were identified using thresholds of |log₂(fold change)| >1, and TPM ≥1 in the higher-expressing cell population.

#### Batch effect correction

To address potential batch effects arising from samples processed at different times, we performed batch effect correction using a limma-style approach. Raw TPM (transcripts per million) values were first filtered to remove low-expression genes (mean TPM < 1 across all samples). The filtered data were then log2-transformed (log2(TPM + 0.01)) to stabilize variance and normalize the distribution.

Batch effect removal was performed in log2 space using a linear model approach equivalent to the removeBatchEffect function from the R limma package.^64^ Genes with zero variance after batch correction were removed from downstream analyses.

### Whole-genome bisulfite sequencing (WGBS)

#### Library preparation and sequencing

Genomic DNA was extracted using the DNeasy Blood & Tissue Kit (Qiagen, Hilden, Germany). Library preparation and sequencing were performed by Rhelixa Japan Inc (Tokyo, Japan). Bisulfite conversion was performed using the EZ DNA Methylation-Gold Kit (ZYMO RESEARCH, Irvine, CA, USA). The eluted DNA was processed using the Accel-NGS Methyl-Seq DNA Library Kit (IDT, Coralville, IA, USA). Libraries were sequenced on an Illumina NovaSeq 6000 platform, generating 150-base paired-end reads.

#### Data analysis

Raw sequencing reads were trimmed using Trim Galore with the recommended conditions for the Accel-NGS Methyl-Seq DNA Library Kit and then aligned to the human reference genome (GRCh38/hg38) using Bismark with default parameters. Bismark eliminated duplicated reads, and methylation calls were made using the Bismark methylation extractor with a minimum sequencing depth of 4 at CpG sites. CpG coverage against GRCh38 was 90% for REC and SrEC, and 89% for bulk MSC. Differentially methylated regions (DMRs) were identified using the R package DSS (≥10% difference of CpG methylation, local FDR threshold = 0.02, minCG = 15, and minlen = 300). Average CpG methylation levels in each enhancer were calculated using bedtools. Only autosomal chromosomes were considered for DNA methylation analysis.

#### Characterization of differentially methylated regions

BED files for promoter- and enhancer-like signatures were downloaded from SCREEN (ENCODE3). Promoter-associated and enhancer-associated DMRs were identified by intersecting with these signatures using bedtools. DMRs within gene bodies were identified by intersecting nonpromoter-associated DMRs with GENCODE V45 genes. DMRs overlapping histone modifications in hESC H1 were identified using bedtools and deeptools.

### Gene Ontology analysis of DEGs and DMR-associated DEGs

To investigate the relationship between enhancer methylation and gene expression, GREAT was used to identify genes within 200 kb (Figure 4) or 1000 kb (Figure 3) of DMR-associated poised enhancers, employing the “single nearest gene” association strategy. These genes were analyzed for differential expression between tREC^L6^ and SrEC^L12^ (>2-fold change) to identify DMR-associated DEGs. GO analysis of the resulting gene sets was performed using GREAT or Metascape with Benjamini-Hochberg correction for multiple testing.

### Statistical analysis of DMR characteristics

Statistical analyses were performed using R software (version 4.3.0) with standard packages. For enrichment analysis of DMRs in genomic features, chi-square tests of independence were conducted using the chisq.test() function in base R. Shuffled control bed files were generated as null distributions for statistical analysis, in which DMR locations were randomly shuffled on the same chromosome using the bedtools “shuffle” tool (v2.30.0) with the -chrom parameter to maintain chromosomal distribution. Overlaps with promoter-like or enhancer-like signatures and gene bodies were recalculated for shuffled controls using bedtools intersect. Chi-square tests of independence were performed to assess whether the observed overlap frequencies significantly differed from expected frequencies based on shuffled controls. Statistical significance was determined with significance levels set at ***P < 1×10^-10^ (highly significant), **P < 1×10^-4^ (significant), and *P < 0.05 (marginally significant). All statistical tests included sufficient sample sizes (minimum n > 1000 DMRs per group) to ensure adequate statistical power, with exact sample sizes indicated in Figure legends. Prior to chi-square testing, expected frequencies in all cells of contingency tables exceeding 5 were verified, satisfying the fundamental assumption for chi-square tests of independence.

### CUT&Tag assay and analysis

#### Assay procedure

The CUT&Tag assay was performed using the CUT&Tag-IT® Assay Kit (Active Motif, Carlsbad, CA, USA), anti-rabbit (#53160, Active Motif), histone H3K4me3 antibody (#39060, Active Motif), and histone H3K27me3 antibody (#39157, Active Motif) according to the manufacturer’s instructions, with slight modifications. Cells attached to a polystyrene dish were washed with chilled PBS twice and covered with ice-cold NE1 buffer for 10 min on ice, then collected with a cell scraper into a 1.5-mL centrifugation tube. Library preparation, and sequencing were performed by Genome-Lead Co., Ltd. Libraries were sequenced on an Illumina NovaSeq 6000 platform, generating 150-base paired-end reads.

#### Data analysis

Raw sequencing reads were trimmed using Trim Galore! (ver. 0.6.4) with default parameters and aligned to the human reference genome (GRCh38/hg38) using Bowtie2 (ver. 2.3.5) with default parameters. Only uniquely mapped reads were selected for further analysis. To establish a baseline reference for the poised enhancer landscape, CUT&Tag for H3K27me3 was additionally performed on commercially available, uncultured bulk human bone marrow-derived MSCs (Cat#PT-2501, Lonza Japan). To ensure comparability, these bulk MSCs were thawed, expanded under culture conditions identical to those used for REC isolation, and harvested at an early passage upon reaching subconfluency. Enhancer-like signature regions overlapping with the H3K27me3 histone modification within a 2-kb window were defined as poised enhancers. Genome tracks were created using pygenometracks (ver. 3.8) to visualize CpG-specific methylation levels and histone ChIP-seq peaks. Heatmaps of histone marks were created using deeptools (ver. 2.0).

### Principal component analysis for poised enhancer-related genes

Autosomal genes were identified from RNA-seq data, excluding those that were not expressed. TPM values were log2-transformed using a pseudocount of 0.01, the minimum non-zero value in the dataset, to enable computation (log2(TPM + 0.01)), and batch effects were corrected using the removeBatchEffect function from the R package limma. To preserve the biological distinction between unexpressed and expressed genes, values corresponding to originally unexpressed genes (TPM = 0) were reset to zero after batch correction. Genes with no expression variation across samples were then removed before conducting PCA. For comparative analysis, PCA was performed on both the poised enhancer-associated gene set (n = 12 968) and the complete transcriptome (n = 54 700) to evaluate the discriminatory power of targeted gene selection compared with whole transcriptome approaches.

### Gene set enrichment analysis (GSEA)

To elucidate the biological functions associated with the principal components, GSEA was performed on the full ranked list of poised enhancer-associated genes. Genes were ranked separately by their loading scores for PC1, and PC2. The analysis was conducted using the prerank module of the GSEApy Python library (version 1.0.3)^65^ against the MSigDB Hallmark 2020 gene set collection (MSigDB_Hallmark_2020).^66^ The analysis was run with 1000 permutations, and gene sets with sizes between 15 and 500 genes were included. Pathways with a FDR q-value < 0.25 were considered significantly enriched.

#### Use of artificial intelligence tools

During the preparation of this manuscript, Claude (Anthropic) and Gemini (Google) to improve the clarity, conciseness, and logical flow of the English text. After using these tools/services, the authors reviewed, and edited the manuscript as needed and take full responsibility for the content of the published article.

## Acknowledgements

This work was supported by a grant from PuREC Co., Ltd. through a collaborative research project with H.K.

We thank past and present lab (Ehime University) and PuREC members, especially Yoshikazu Morishita and Erika Miwa, for providing valuable insights and discussion. We would like to acknowledge Editage (www.editage.com) for English language editing.

## Author contributions

Data collection was conducted collaboratively by PuREC Co. Ltd. staff (A.W. and T.S.) and Ehime University researchers (K.H-K. and H.K.). All data analyses were performed exclusively by K.H-K. and H.K. from Ehime University. This strict division of responsibilities ensured analytical independence and scientific integrity in interpreting the results. H.K. and A.W. conceptualized the study, while K.H-K. and A.W. developed the methodology. K.H-K., A.W., and T.S. conducted the investigation, and K.H-K. handled data visualization. H.K. and K.H-K. wrote the original draft, which was reviewed and edited by H.K., K.H-K., A.W., T.S., and Y.M.

---

**Supplementary figure S1.**
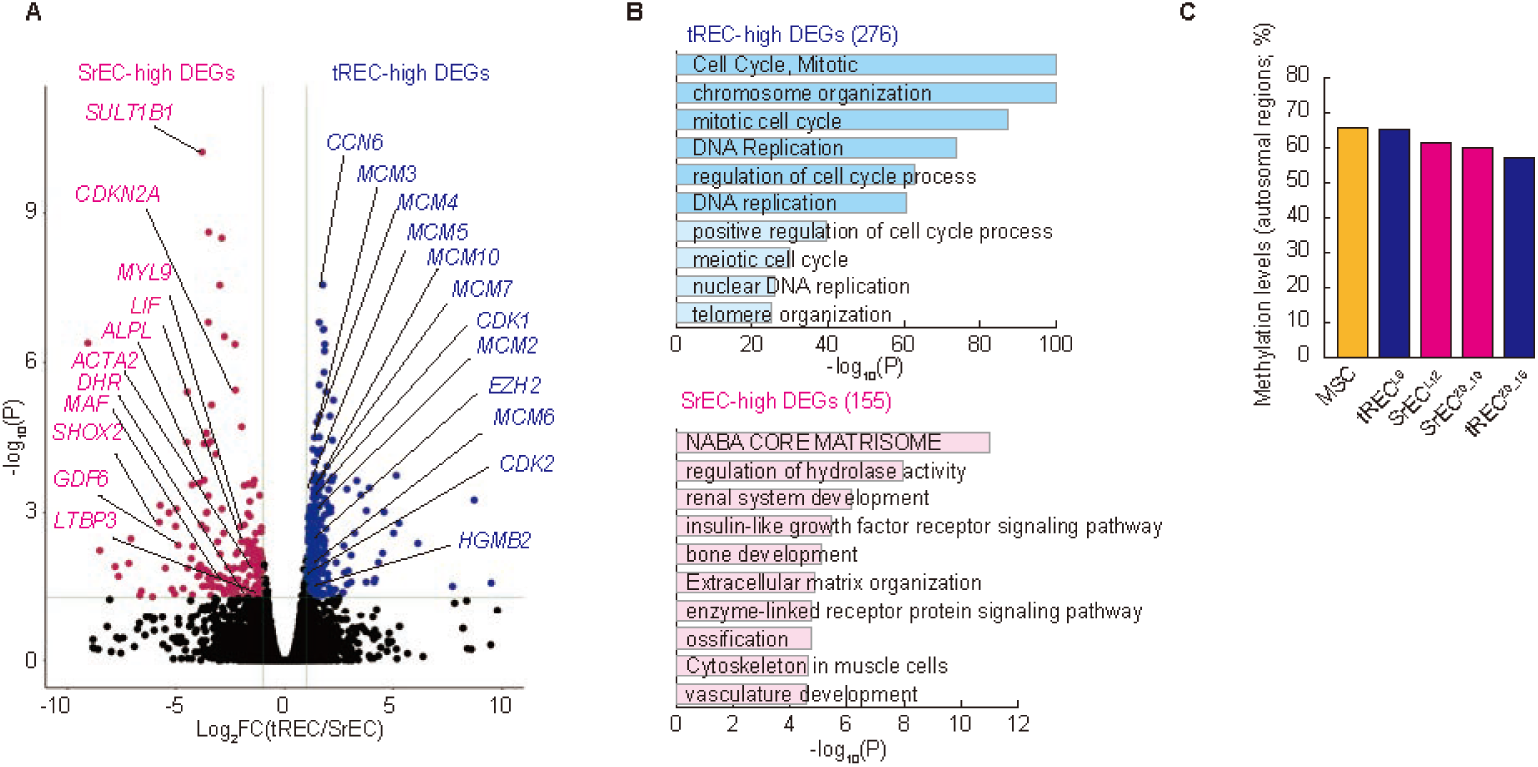
Global transcriptomic and epigenetic profiles of tRECs and SrECs. (A) Volcano plot illustrating the widespread transcriptional differences between the combined tREC (blue dots) and SrEC (red dots) populations. Key genes associated with the tREC (e.g., *EZH2*, *MCM* family) or SrEC (e.g., *CDKN2A*, *ALPL*) states are highlighted. (B) GO analysis of differentially expressed genes. tRECs are enriched for cell cycle and proliferation pathways (e.g., cell cycle, mitotic, DNA replication), whereas SrECs are enriched for pathways related to the matrisome (e.g., NABA CORE MATRISOME), and lineage-specific development (e.g., bone development, ossification). (C) Global CpG methylation levels (autosomal regions) as determined using WGBS. Contrary to the global hypomethylation often associated with extensive cell culture, the elite tREC^L6^ clone maintains a high level of global methylation, comparable to that in the bulk MSC population and higher than that in its isogenic SrEC^L12^ counterpart. Abbreviations: GO, Gene Ontology; MSC, mesenchymal stem cells; tRECs, “true” rapidly expanding clones; SrECs, subrapidly expanding clones; WGBS, Whole genome bisulfite sequencing

**Supplementary figure S2.**
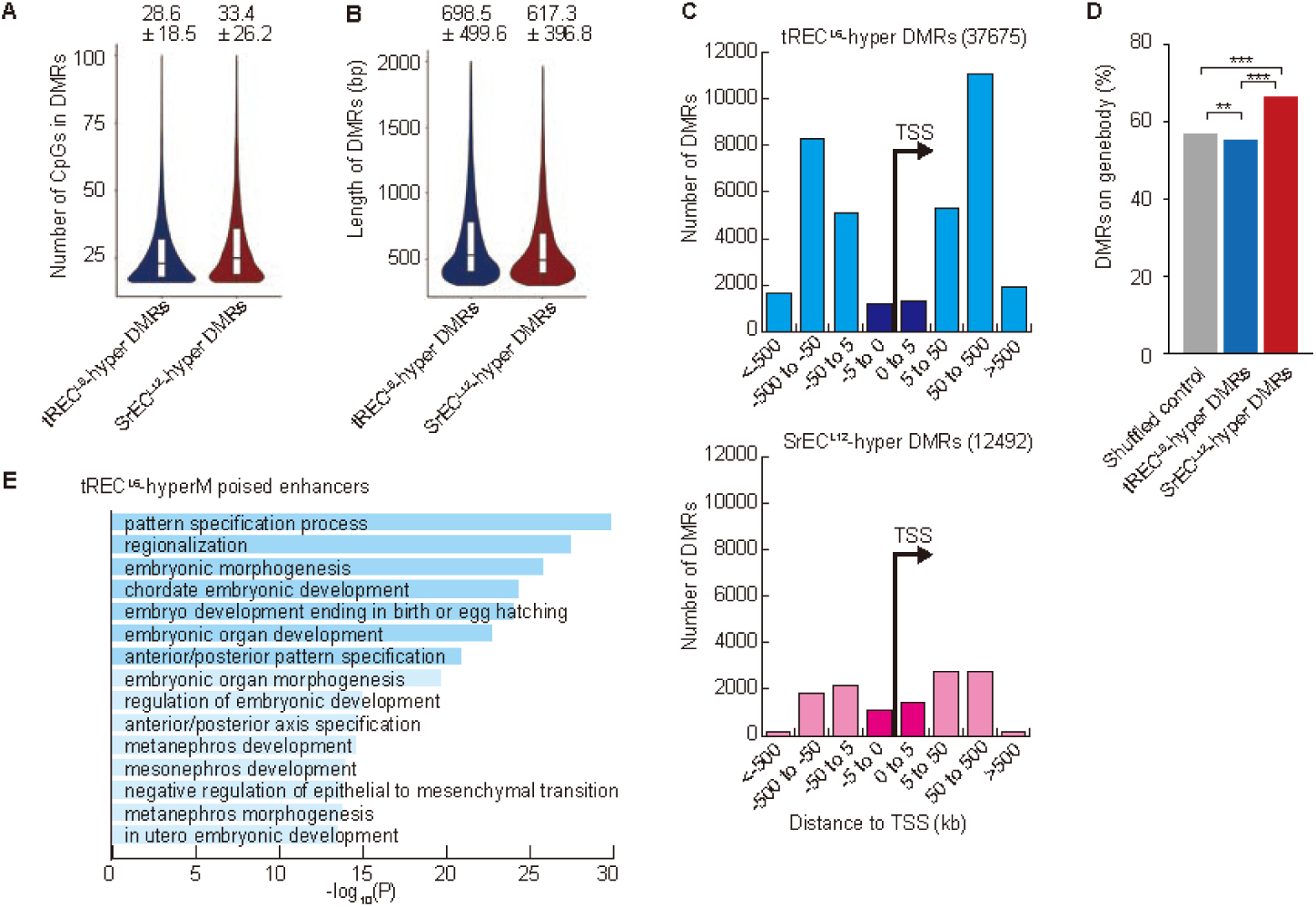
Distinct characteristics of hypermethylated DMRs in tREC^L6^ and SrEC^L12^. (A and B) Violin plots comparing the properties of tREC^L6^-hyperM and SrEC^L12^-hyperM DMRs. SrEC^L12^-hyperM DMRs are characterized by higher CpG density (A) and smaller size (B) compared with tREC^L6^-hyperM DMRs, suggesting CpG island targeting(Wilcoxon signed-rank test, p < 2.2 × 10^-16^). (C and D) Genomic distribution of DMRs relative to gene structures. In stark contrast to tREC^L6^-hyperM DMRs, which are broadly scattered, SrEC^L12^-hyperM DMRs are significantly enriched near TSSs (C) and within gene bodies (D) (Data represent comparison vs. shuffled control). (E) GO analysis of genes associated with tREC^L6^-hypermethylated poised enhancers. Although this enrichment comprises broad, “upstream” developmental terms (e.g., pattern specification process, embryonic morphogenesis) reflecting the inherent nature of poised enhancers, it notably lacks the specific enrichment for downstream MSC lineages, such as skeletal system development, which is characteristic of the SrEC^L12^ profile (cf. Figure 3E). Abbreviations: DMR, differentially methylated regions; GO, Gene Ontology; MSC, mesenchymal stem cells; tRECs, “true” rapidly expanding clones; SrECs, subrapidly expanding clones; TSS, transcription start site

**Supplementary figure S3.**
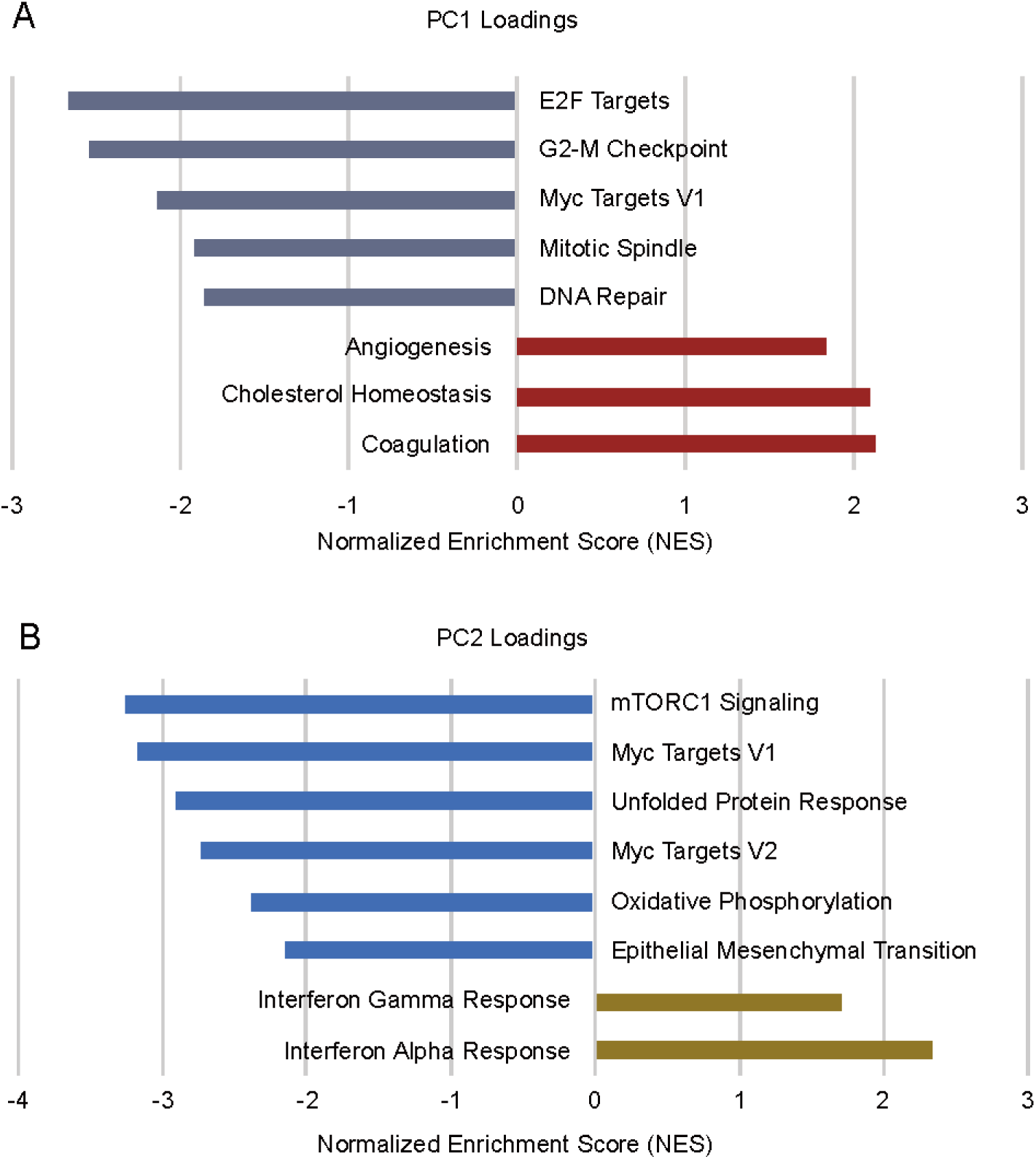
GSEA confirms the biological identity of the principal components. GSEA was performed on the ranked list of genes contributing to PC1 and PC2 from the poised enhancer-associated gene set. (A) PC1 loadings: This analysis confirms the biological identity of PC1 as the primary axis of functional quality.

- Negative loadings (characteristic of the high-quality tREC state) are strongly enriched for pathways essential for proliferation and genome maintenance (e.g., E2F targets, G2-M checkpoint, DNA repair).
- Positive loadings (characteristic of the low-quality SrEC state) are enriched for pathways such as angiogenesis and coagulation. Critically, inspection reveals that the “coagulation” gene set is driven by core mesenchymal differentiation factors, supporting our model in which PC1 represents a biological axis from a multipotent (tREC) to a prodifferentiated (SrEC) state. (B) PC2 loadings: This analysis provides a powerful biological interpretation for the PC2 axis. PC2 loadings are strongly enriched for pathways reflecting individual immune status, such as interferon gamma response and interferon alpha response. This finding provides a mechanistic basis suggesting that PC2 effectively segregates clones by their donor origin, for instance, separating the distinct clusters derived from donor BM lot# 082064B (e.g., tREC^L6^, SrEC^L12^, Clone H) and donor BM lot# 18TL352886 (e.g., tREC^20_15^, SrEC^20_10^, Clone E), as shown in Figure 5A (see also Supplementary Table S1). Abbreviations: GSEA, Gene set enrichment analysis; tRECs, “true” rapidly expanding clones; SrECs, subrapidly expanding clones

**Supplementary figure S4.**
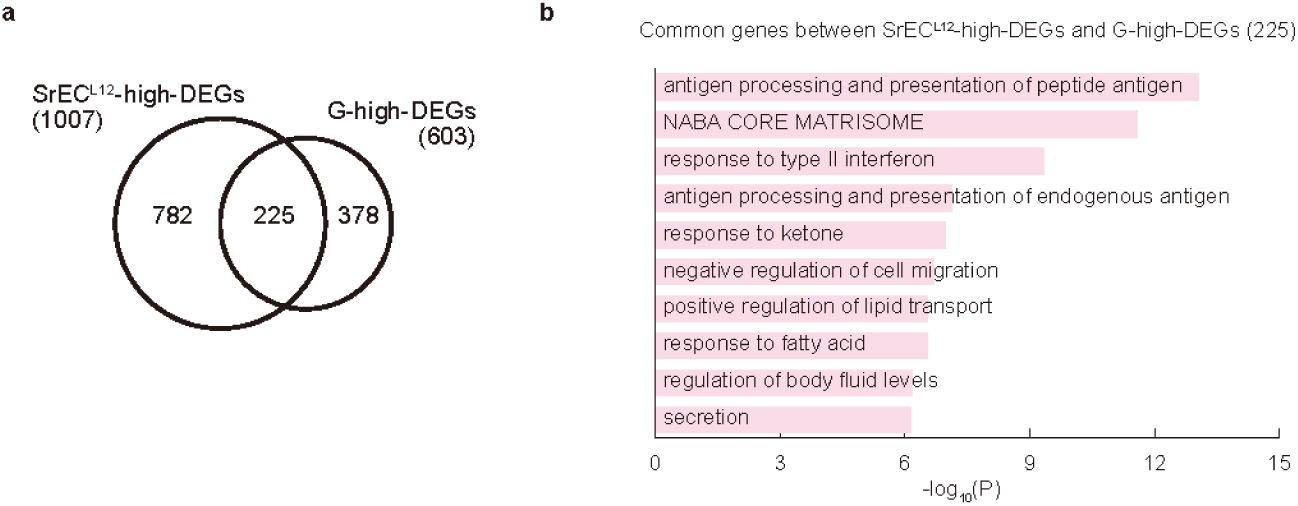
Candidate clone G shares a conserved transcriptional signature of functional decline with SrECs. (A) Venn diagram showing the significant overlap (225 genes) between genes upregulated in the reference SrEC^L12^ (1007 genes total) and the prospectively classified SrEC-like clone G (603 genes total). (B) GO analysis of the 225 commonly upregulated genes. This shared signature is highly characteristic of the SrEC phenotype, showing enrichment for pathways such as antigen processing and presentation of peptide antigen, NABA CORE MATRISOME, and response to type II interferon. Abbreviations: GO, Gene Ontology; MSC, mesenchymal stem cells; SrECs, subrapidly expanding clones; PCA, principal component analysis

**Supplementary figure S5.**
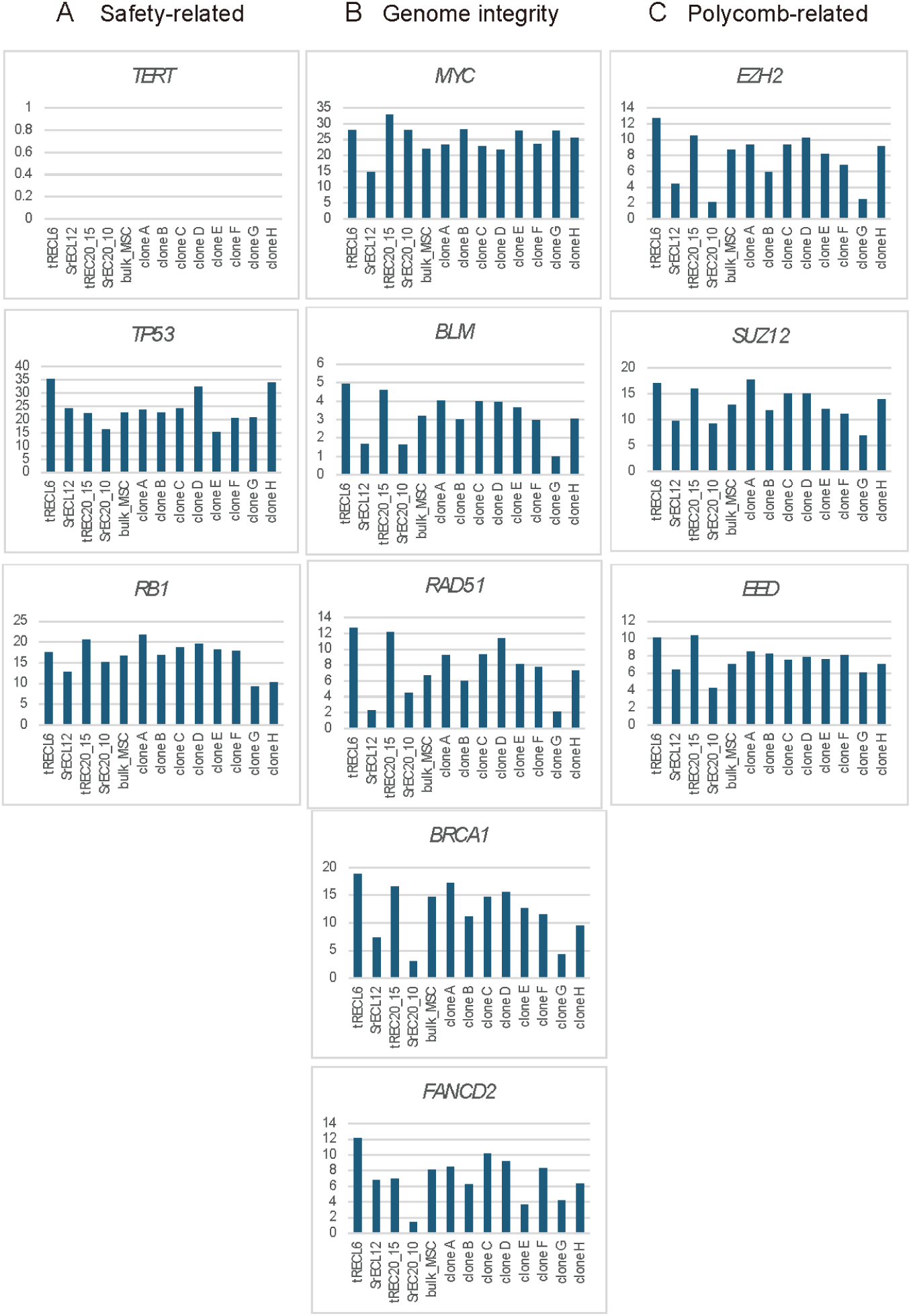
tRECs are characterized by intact safety checkpoints and robust genome and epigenome maintenance networks. Bar charts showing the expression levels (TPM) of key gene networks across all MSC clones. (A) Safety-related: All clones are negative for *TERT* expression, confirming that they are not immortalized. Superior tREC-like clones (e.g., tREC^L6^, tREC^20_15^, Clone A) maintain robust expression of the key tumor suppressors *TP53* and *RB1*. (B) Genome integrity: tRECs show high expression of the “genomic guardian” network, including key DNA repair and stability genes, such as *MYC*, *BLM*, *RAD51*, *BRCA1*, and *FANCD2*. (C) Polycomb-related: The core components of the Polycomb repressive complex 2 (PRC2), namely *EZH2*, *SUZ12*, and *EED*, are coordinately upregulated in tRECs. This finding is consistent with their role as “epigenomic guardians” that maintain the poised enhancer landscape, a key mechanism proposed in the present study. Abbreviations: MSC, mesenchymal stem cells; tRECs, “true” rapidly expanding clones; TPM, transcripts per million

## SUPPLEMENTARY INFORMATION

Tables S1–S3

## SUPPLEMENTARY TABLES

**Supplementary Table S1.** Characteristics of MSC clones used in the present study. All MSC clones used in the present study are summarized. This table details the functional classification (tREC or SrEC), donor of origin (BM lot#), and other donor characteristics (age, sex, race). The BM lot# column is critical for identifying isogenic clone pairs (e.g., tRECᴸ⁶/SrEC^L12^) and assessing the donor-specific effects (e.g., Figure 5A and Supplementary Figure S5B). The passage 3 proliferation index (P3-PI) is also defined and the adipogenic and osteogenic differentiation potential is confirmed.

|  | REC lot# | BM lot# | Age | Sex | Race | P3-PI | Adipogenesis | Osteogenesis |
| --- | --- | --- | --- | --- | --- | --- | --- | --- |
| tREC<sup>L6</sup> | 2209L#6 | 082064B | 23 | Male | Black or African American | 10 | Positive | Positive |
| SrEC<sup>L12</sup> | 2209L#12 | 082064B | 23 | Male | Black or African American | 6.4 | Positive | Positive |
| tREC<sup>20_15</sup> | 2010#20-15 | 18TL352886 | 23 | Male | Black or African American | 24.4 | Positive | Positive |
| SrEC<sup>20_10</sup> | 2010#20-10 | 18TL352886 | 23 | Male | Black or African American | 17.9 | Positive | Positive |
| MSC | ND (Lonza) | 2M-125C | ND | Male | Black or African American | ND | ND | ND |
| Clone A | 1904fBM#3 | 3021747 | 23 | Male | Black or African American | 23.6 | Positive | Positive |
| Clone B | 1707(1)#4 | 503704 | 24 | Male | Black or African American | 18.5 | Positive | Positive |
| Clone C | 1911#10 | 082064B | 23 | Male | Black or African American | 10.6 | Positive | Positive |
| Clone D | 1911#13 | 082064B | 23 | Male | Black or African American | 11.9 | Positive | Positive |
| Clone E | 2010#20-13 | 18TL352886 | 23 | Male | Black or African American | 32.1 | Positive | Positive |
| Clone F | 1708#3 | BM6845 | 22 | Male | Hispanic | 14.5 | Positive | Positive |
| Clone G | 1807#5 | 503704 | 24 | Male | Black or African American | 25 | Positive | Positive |
| Clone H | 1602#6 | 082064B | 23 | Male | Black or African American | 12.7 | Positive | No Data |

Abbreviations: MSC, mesenchymal stem cell; tRECs, “true” rapidly expanding clones; SrECs, subrapidly expanding clones; BM, bone marrow; ND, not determined (or not disclosed by the manufacturer).

**Supplementary Table S2.** Expression of transcription factors whose binding motifs are enriched at SrEC-hypermethylated poised enhancers. Batch-corrected TPM values are listed for the key transcription factors whose binding motifs were identified using HOMER analysis in Figure 3G. The data confirm the robust expression of these regulators (e.g., *RUNX2*, *SMAD3*, *TEAD1*) across all analyzed MSCs, including the SrEC clones. This finding fulfills a key prerequisite for the “poised enhancer decommissioning” mechanism, demonstrating that the transcriptional “keys” (TFs) required to activate these newly accessible (hypermethylated) enhancers are readily available.

|  | tREC<sup>L6</sup> | SrEC<sup>L12</sup> | tREC<sup>20_15</sup> | SrEC<sup>20_10</sup> | Bulk MSC |
| --- | --- | --- | --- | --- | --- |
| TEAD1 | 79.62 | 59.49 | 57.87 | 57.97 | 61.22 |
| SMAD3 | 15.23 | 23.24 | 26.35 | 25.09 | 27.42 |
| RUNX2 | 6.05 | 3.01 | 19.97 | 15.20 | 15.08 |
| MEF2A | 14.96 | 37.33 | 11.83 | 11.91 | 20.37 |
| TEAD4 | 8.65 | 5.09 | 7.47 | 3.40 | 6.98 |
| FLI1 | 3.38 | 3.20 | 6.40 | 5.05 | 4.10 |
| HIF1A | 130.55 | 101.90 | 282.58 | 319.11 | 243.91 |
| JUNB | 11.73 | 21.24 | 21.61 | 15.25 | 21.08 |
| CTCF | 18.32 | 12.21 | 16.92 | 9.69 | 14.97 |

Abbreviations: MSC, mesenchymal stem cell; SrECs, subrapidly expanding clones; TPM, transcripts per million

**Supplementary Table S3.**
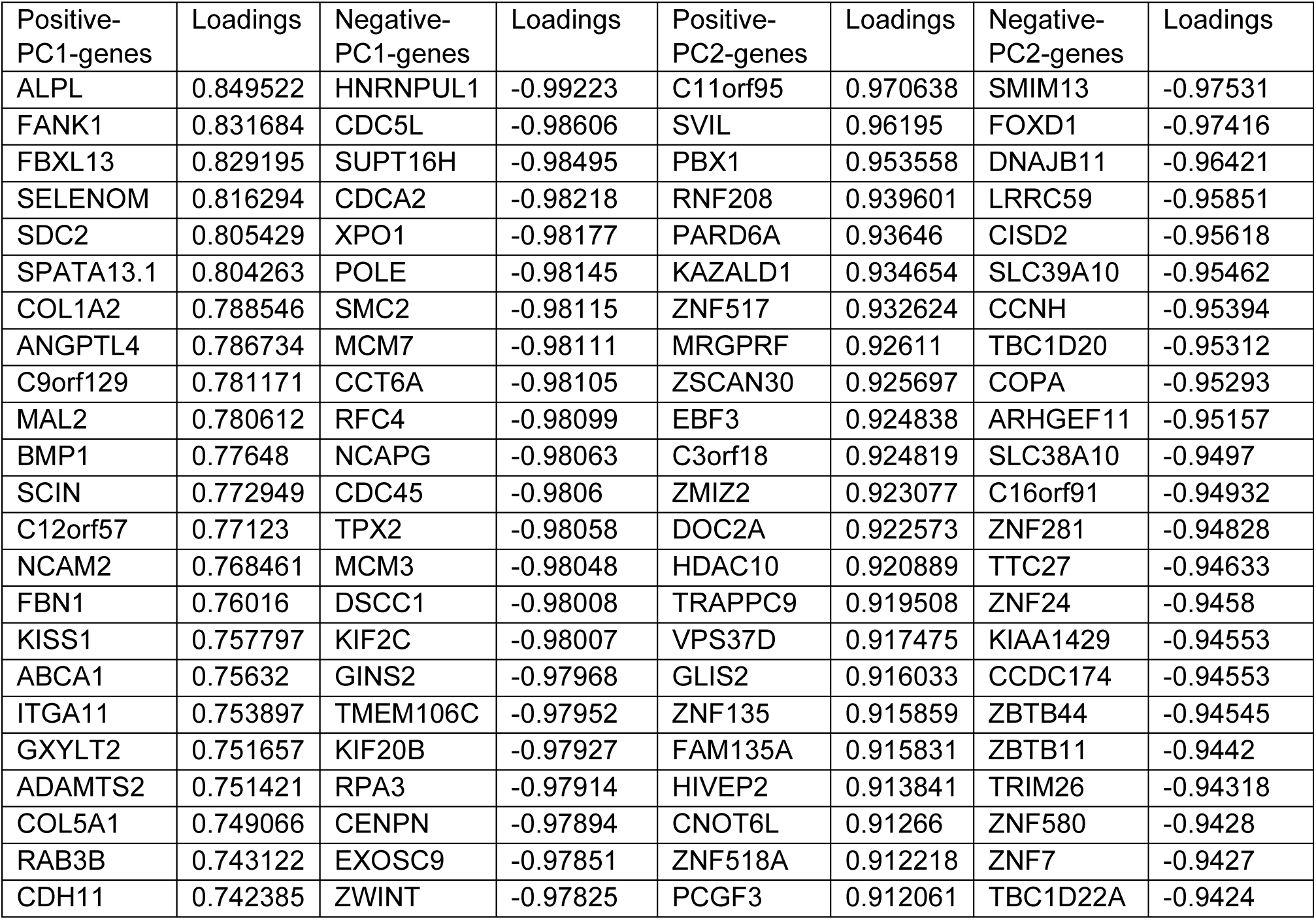
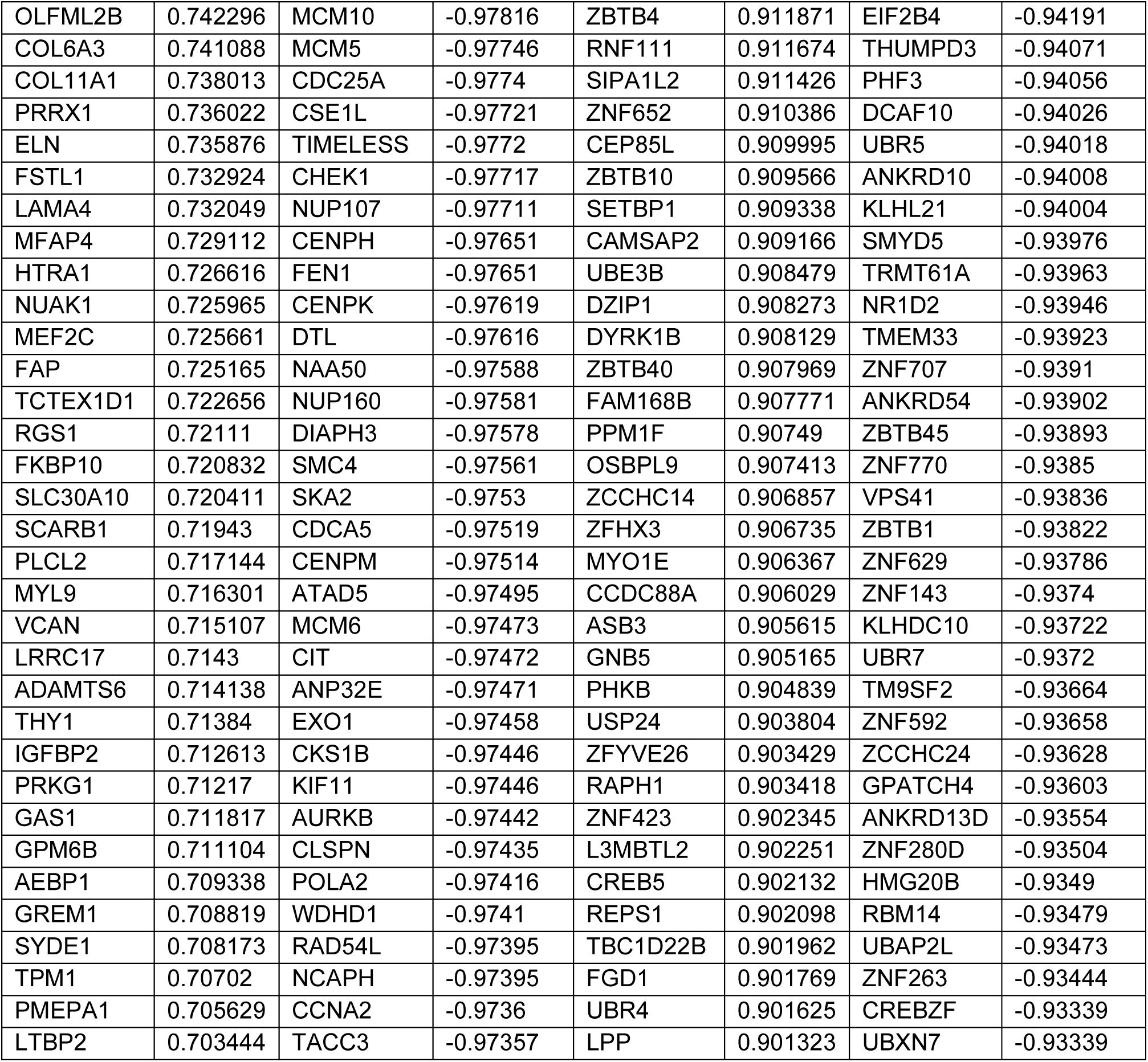
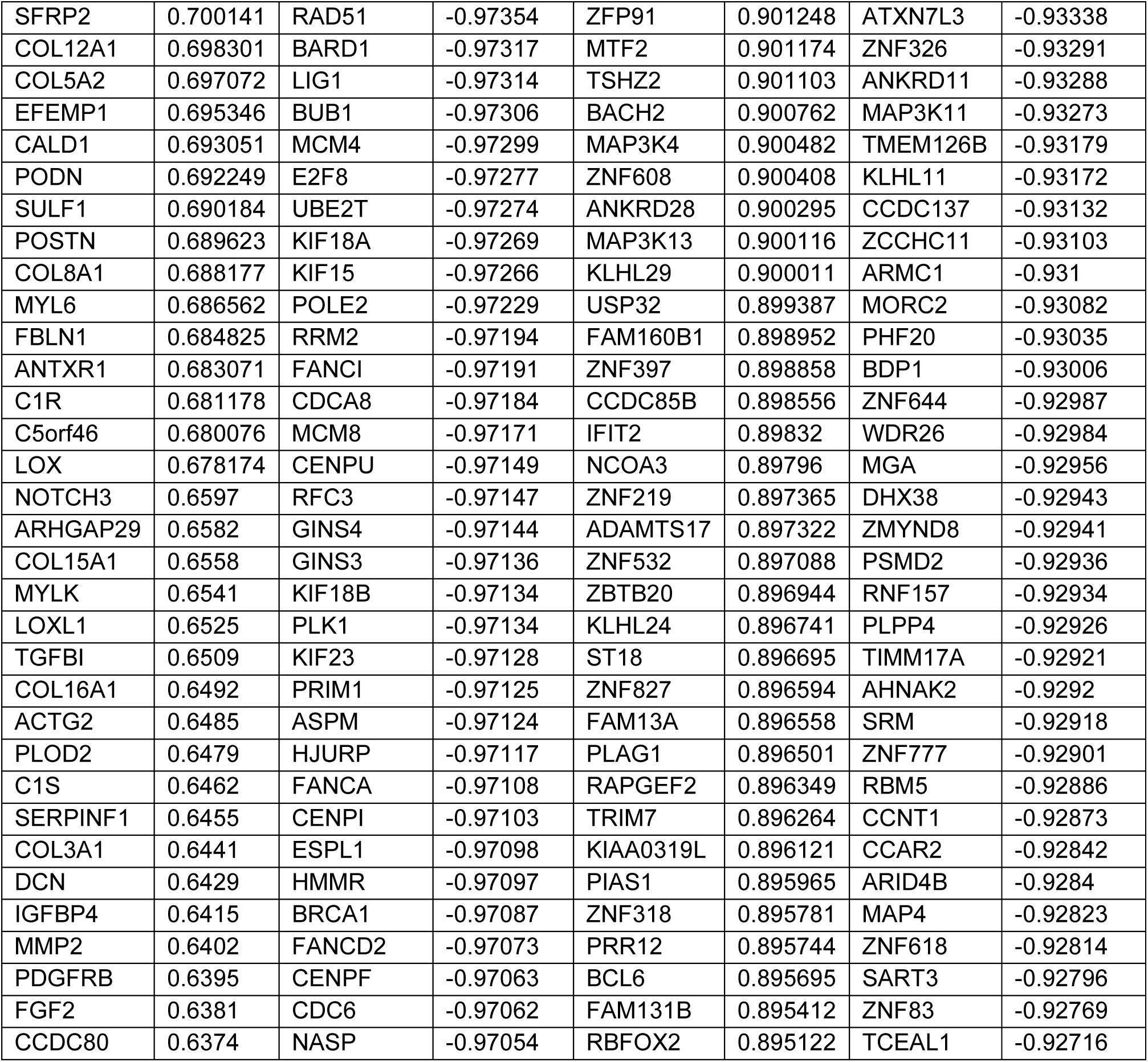
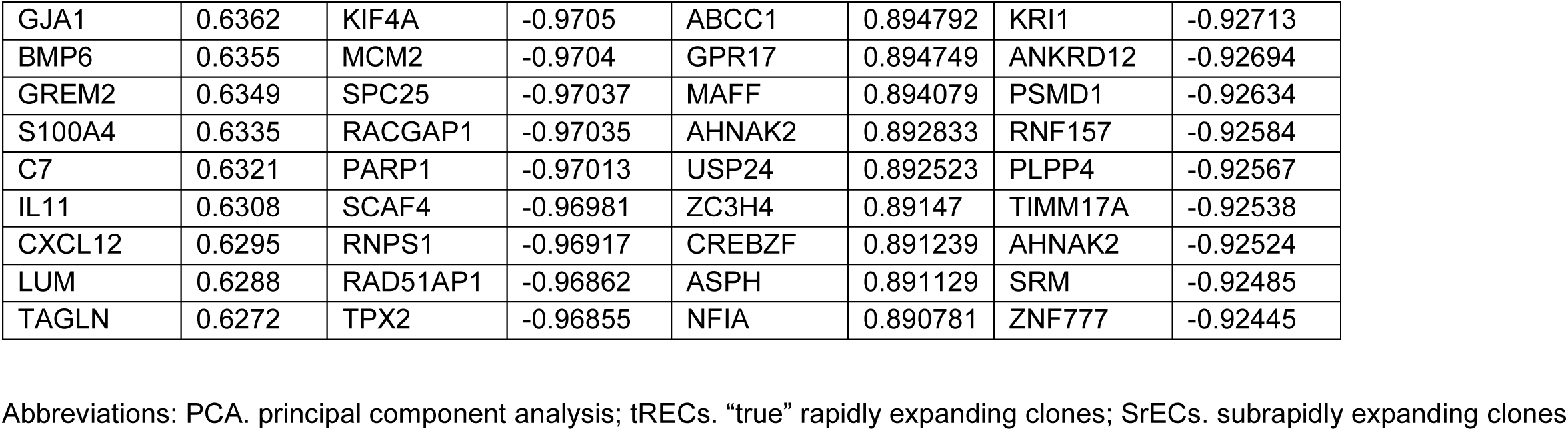
Gene loadings for principal components 1 and 2 from PCA of poised enhancer-associated genes. This table lists the loading scores for the top 100 positively and negatively contributing poised enhancer-associated genes on the first two principal components (PC1 and PC2), derived from the analysis shown in Figure 5A and interpreted in Supplementary Figure S5. PC1 loadings represent the contribution of each gene to the primary axis of functional quality. Genes with high positive scores (e.g., *ALPL*) drive the inferior SrEC state, whereas genes with high negative scores drive the superior tREC state (enriched for proliferation pathways, see Supplementary Figure S5A). PC2 loadings represent the contribution to the secondary axis of variation, which captures the donor-specific physiological signatures (e.g., interferon response), as interpreted in Supplementary Figure S5B.

